# Triploidy is prominent in the duckweed *Lemna minor* complex

**DOI:** 10.1101/2025.02.18.638736

**Authors:** Todd P. Michael, Phuong T.N. Hoang, Bradley W. Abramson, Allen Mamerto, Buntora Pasaribu, Evan Ernst, Nicholas Allsing, Luca Braglia, Semar Petrus, Jörg Fuchs, Veit Schubert, Nolan Hartwick, Megan Wang, Mariele Lensink, Tiffany Duong, Kelly Colt, Manuela Bog, K. Sowjanya Sree, Laura Morello, Klaus-J. Appenroth, Ingo Schubert, Robert A. Martienssen, Eric Lam

**Affiliations:** The Plant Molecular and Cellular Laboratory, Salk Institute for Biological Studies, La Jolla, CA 92037 USA; Department of Cell and Developmental Biology, School of Biological Sciences, and Center for Marine Biotechnology and Biomedicine, University of California, San Diego, La Jolla, CA, 92093, USA; San Diego Botanical Garden, Encinitas, CA, 92024, USA; Leibniz Institute of Plant Genetics and Crop Plant Research (IPK) Gatersleben, D-06466 Seeland, Germany; Biology Faculty, Dalat University, District 8, Dalat City, Lamdong Province, Vietnam; Department of Plant Biology, Rutgers the State University of New Jersey, New Brunswick, NJ 08901 USA; Howard Hughes Medical Institute, Cold Spring Harbor Laboratory, Cold Spring Harbor, NY, USA; Institute of Agricultural Biology and Biotechnology, National Research Council (CNR), Milan, Italy; Institute of Botany and Landscape Ecology, University of Greifswald, D-17489 Greifswald, Germany; School of Biotechnology, Institute of Science, Banaras Hindu University, Varanasi-221005, India; Matthias Schleiden Institute-Plant Physiology, University of Jena, D-07743 Jena, Germany

**Keywords:** Genome sequencing, *Lemna minor*, genome plasticity, genome size, genomic *in situ* hybridization, interspecific hybridization, super-pangenome, pangenome, ploidy, triploidy

## Abstract

Duckweeds (Lemnaceae Martinov) are aquatic monocotyledonous flowering plants comprising five genera and 35 recognized species, known for being the smallest and fastest-growing flowering plants on Earth. Many species are morphologically indistinguishable due to their highly reduced structures, yet molecular evidence suggests that visually similar clones may represent distinct species or hybrids. For example, clonal accessions of the globally distributed *Lemna minor* in the Landolt Duckweed Collection exhibit genome size variations of several hundred megabases (Mb), raising questions about their taxonomic classification and evolutionary origins. We analyzed 58 presumed *L. minor* clones to resolve these relationships using a comprehensive suite of methods, including whole-genome sequencing (WGS), flow-cytometric genome size measurements, molecular markers, chromosome counting, and genomic *in situ* hybridization (GISH). Our findings reveal extensive genome plasticity within the “*Lemna minor* complex,” identifying diploid and triploid *L. minor* clones, as well as di-haploid and triploid interspecific hybrids called *L. × japonica* (*L. minor* × *L. turionifera*), *L. × mediterranea* (*L. minor* × *L. gibba*), and a novel African-clade distinct from known *L. minor* lineages. Triploidy was prevalent, occurring in 29% of the clones, and was associated with enhanced growth under optimal conditions but reduced performance under high light and temperature. These findings highlight the widespread role of triploidy, cryptic species, and hybridization in the *L. minor* complex, emphasizing the importance of multiple approaches for accurately classifying duckweed species and understanding their evolutionary trajectories.

## Introduction

Duckweeds (family Lemnaceae) are rapidly gaining recognition as versatile organisms with broad applications, owing to their simple body plan, rapid growth, and adaptability to various aquatic environments (Lam and Michael, 2022). As emerging model plants, their small genomes and fast life cycles facilitate genetic and physiological research, enabling insights into fundamental biological processes (Acosta *et al*., 2021; Mateo-Elizalde *et al*., 2023). Their high protein content has spurred interest in duckweed as a sustainable feed supplement and a potential human food source, including for bioregenerative life-support systems in space (Hillman and Culley, 1978; Romano and Aronne, 2021; Appenroth *et al*., 2017). Duckweeds are also effective at phytoremediation, removing pollutants and excess nutrients from wastewater, making them ideal for environmental management (Cheng and Stomp, 2009). Beyond these research and ecological applications, the rapid biomass accumulation of duckweed has prompted investigations into industrial uses such as biofuel production and biomaterials development (Liang *et al*., 2023; Guo *et al*., 2023). Altogether, these attributes position duckweed as a promising resource for addressing challenges in food security, environmental sustainability, and innovative plant research on Earth and beyond.

The Duckweed family comprises 35 species and two recognized hybrids across five genera, *Spirodela*, *Landoltia*, *Lemna*, *Wolffia*, and *Wolffiella*, which vary in size from about 15 mm down to less than 1 mm (Acosta *et al*., 2021). *Spirodela*, the most basal genus, features the largest fronds with 10–15 roots, whereas the monospecific *Landoltia* and single-rooted *Lemna* species have smaller fronds and fewer roots. *Wolffia* and *Wolffiella*, the most derived and smallest of the duckweed, lack roots and some *Wolffia* species can double their biomass in under 24 hours (Michael *et al*., 2020). Although duckweeds primarily reproduce asexually by budding daughter fronds from mother fronds, at least some species can also reproduce sexually via flowering and seed formation (Fu *et al*., 2017). Before *Arabidopsis thaliana* became the model plant of choice, duckweeds were central to reductionist plant biology research providing key discoveries in flowering time, circadian rhythms, photosynthesis, and phytohormones (Hillman, 1976). Coupled with its ability to flower, make crosses, be transformed, and thrive under simple conditions, duckweed has become a powerful addition to the model plant toolbox for next-generation molecular studies of plant cells (Acosta *et al*., 2021; Mateo-Elizalde *et al*., 2023).

Since duckweed predominantly grows clonally, and flowering as well as fertile seeds are rare, they must be maintained as a living collection to facilitate researchers working on the same genetic backgrounds. Therefore, Prof. Elias Landolt established and maintained the Landolt Duckweed Collection, now housed at the Rutgers Duckweed Stock Cooperative (RDSC; http://www.ruduckweed.org/), and the National Research Council, CNR, Milan, Italy (https://biomemory.cnr.it/collections/CNR-IBBA-MIDW), which represents together more than 1,000 clones across all five genera and their species collected worldwide by researchers from 1953 to the present (Landolt, 1986). As a first step to characterize this collection, the genome sizes were estimated from selected clones revealing an order of magnitude difference across the family from 160 Mb (*Spirodela polyrhiza*) to 2,203 Mb/1C (*Wolffia arrhiza*) (Wang *et al*., 2011; Van Hoeck *et al*., 2015; Bog *et al*., 2015; Acosta *et al*., 2021; Hoang *et al*., 2022).

Considerable intraspecific genome size variation was noted for some species like *L. minor*, which is known as Common Duckweed since it is the most abundant and cosmopolitan, that ranged from 356-604 Mb yet the differences did not correlate with increased ribosomal DNA or other features tested (Wang *et al*., 2011). The problem with *L. minor* is that it is hard to distinguish morphologically between different accessions and most of the other *Lemna* species (Figure S1) (Volkova *et al*., 2023). Recent molecular and genomic evidence suggests that clones labeled as *L. minor* in the RDSC may represent different species or previously cryptic hybrids such as *L. minor × L. turionifera* (*L.× japonica*) and *L. minor × L. gibba* (*L. × mediterranea*) (Braglia, Lauria, *et al*., 2021; Braglia, Breviario, *et al*., 2021; Schmid *et al*., 2024; Volkova *et al*., 2023; Braglia *et al*., 2024). Therefore, the exact identity for most of the *L. minor* clones in the collection remains ambiguous.

In this work, we systematically evaluated 58 presumed *L. minor* Landolt clones from the RDSC to determine their species identity and genomic traits. Our approach integrated whole-genome sequencing (WGS), flow-cytometric genome size measurements, molecular markers, chromosome counting, and genomic *in situ* hybridization (GISH). We also leveraged our previous chromosome-scale genomes of *L. minor*, *L. gibba*, *L. turionifera*, and *L.× japonica* along with short read (SR) Illumina assemblies to construct a super-pangenome and examine genome dynamics within the “*Lemna minor* complex” (Ernst *et al*., 2023; Todd P. Michael *et al*., 2020; Michael *et al*., 2017; Abramson *et al*., 2021; Baggs *et al*., 2022). We clarified genome composition, ploidy levels, relatedness, and evolutionary origins of the analyzed clones, thereby explaining the observed variability in genome size and chromosome number.

## Results

### Resequencing L. minor from the Landolt Duckweed Collection

We obtained 58 putative *L. minor* clones from the Rutgers Duckweed Stock Cooperative (RDSC) (http://www.ruduckweed.org/) for Illumina short read (SR) resequencing to investigate the genomic variation and potential misidentifications within clones labeled as *Lemna minor*.

Previous studies have reported that some presumed *L. minor* clones in the RDSC may actually be hybrids, specifically *L. minor* × *L. turionifera* (*L. × japonica*) and *L. minor* × *L. gibba* (*L. × mediterranea*) (Braglia, Lauria, *et al*., 2021; Braglia, Breviario, *et al*., 2021; Schmid *et al*., 2024; Volkova *et al*., 2023; Braglia *et al*., 2024; Ernst *et al*., 2023). Additionally, genome size estimates indicate up to two-fold variations among these clones, raising further questions about their classification (Wang *et al*., 2011). Assuming a 1C genome size of ∼400 Mb, we targeted a median sequencing coverage of 60× per haploid genome (Table S1, S2). We mapped the reads to a concatenated reference genome of *L. minor* (Lmin9252) and *L. turionifera* (Ltur9434) to determine hybrid presence and genome composition (Ernst et al., 2023). The overall mapping rate was 98% for all of the clones except four (9425a [67%], 7264 [29%], 5635 [30%], and 8292 [11%]) (Table 1, S1). Recent barcoding studies identified clone 9425a as *L. × mediterranea*, a triploid hybrid between *L. minor* and *L. gibba*, containing one *L. minor* and two *L. gibba* genomes (GGM) (Braglia *et al*., 2024; Romano *et al*., 2024). We remapped clone 9425a to a concatenated version of the *L. minor* (Lmin9252) and *L. gibba* (Lgib7742a) genomes, which increased mapping from 67% to 98% confirming that this clone was *L. × mediterranea*.

**Table 1.**
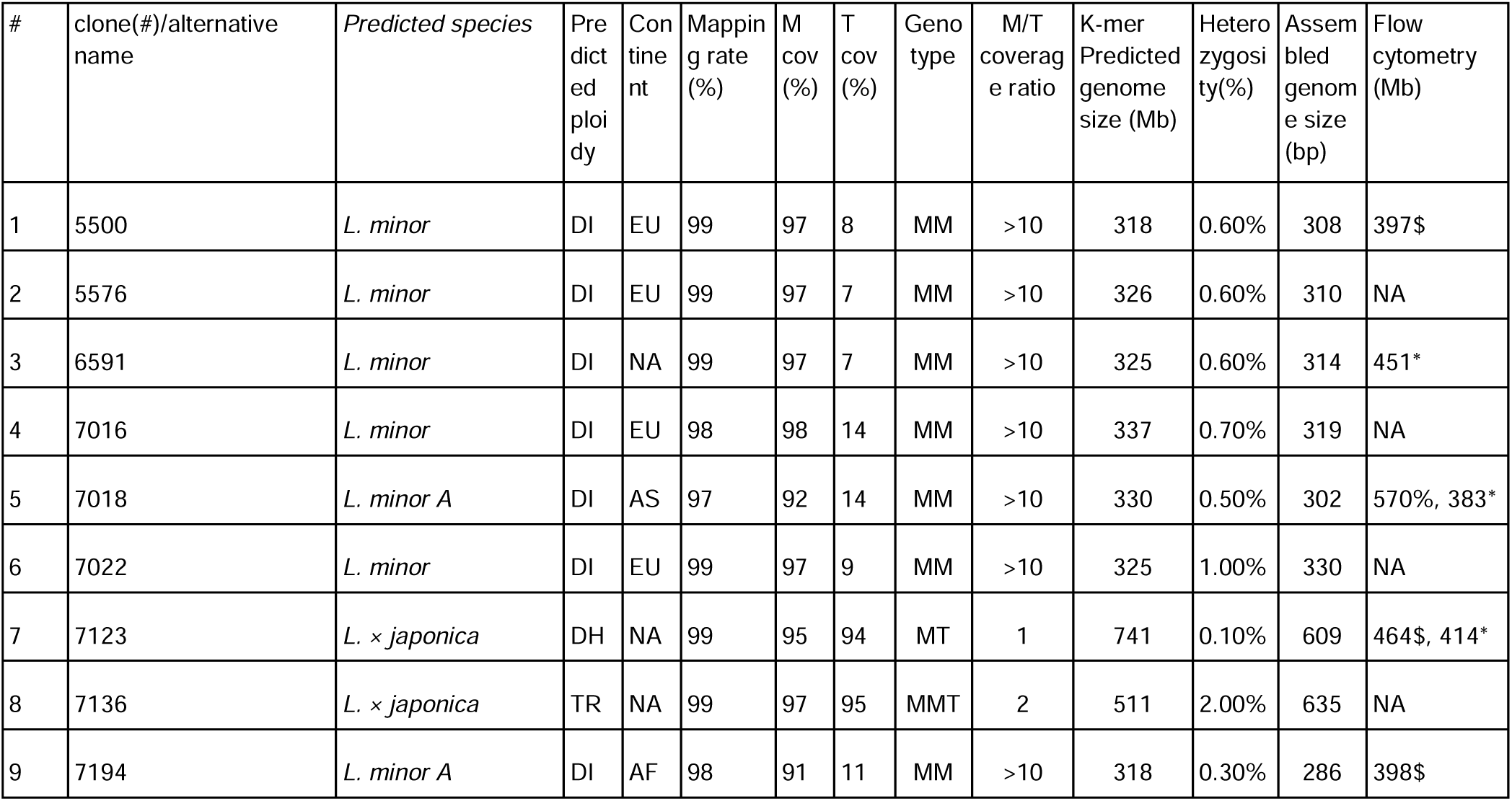
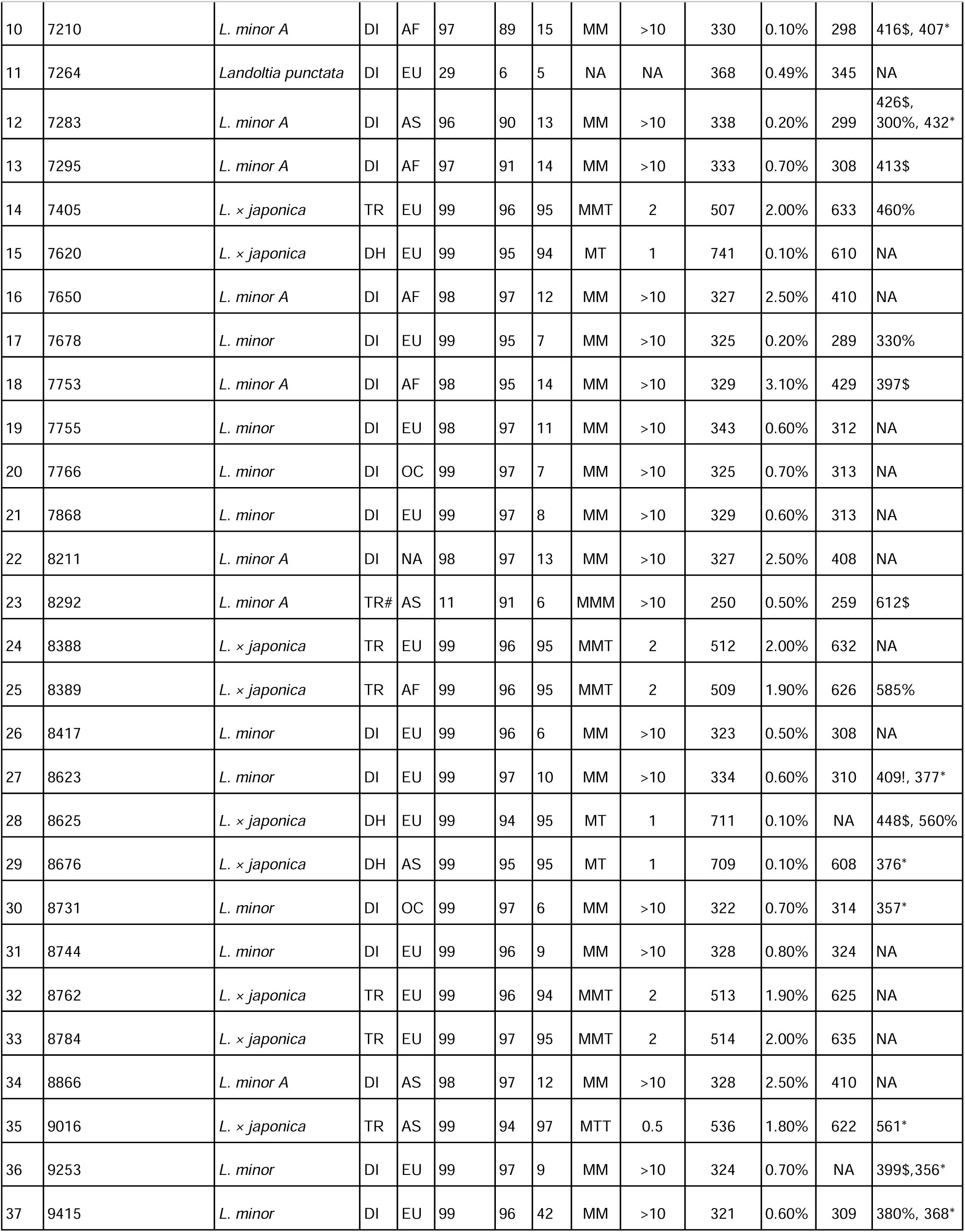
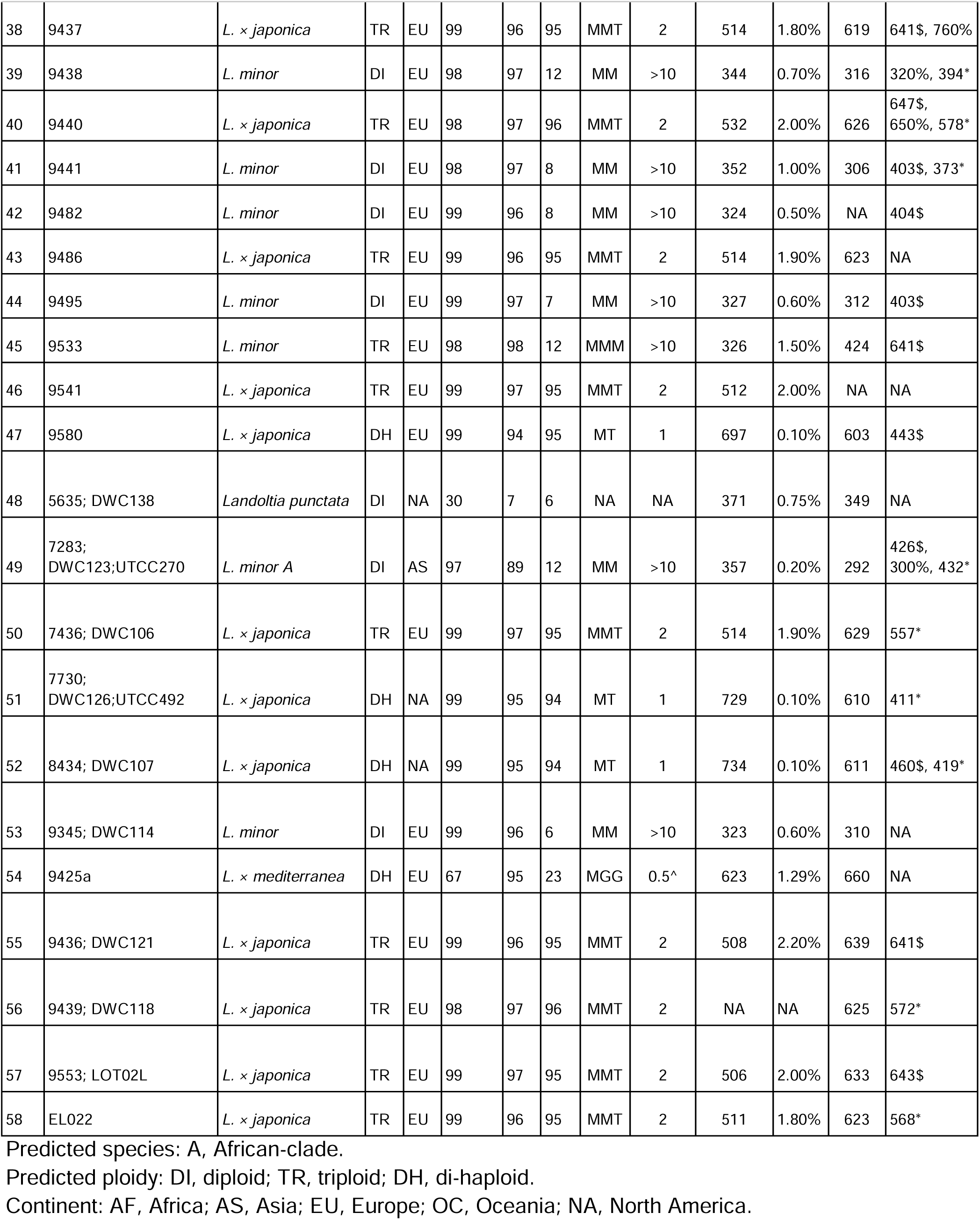
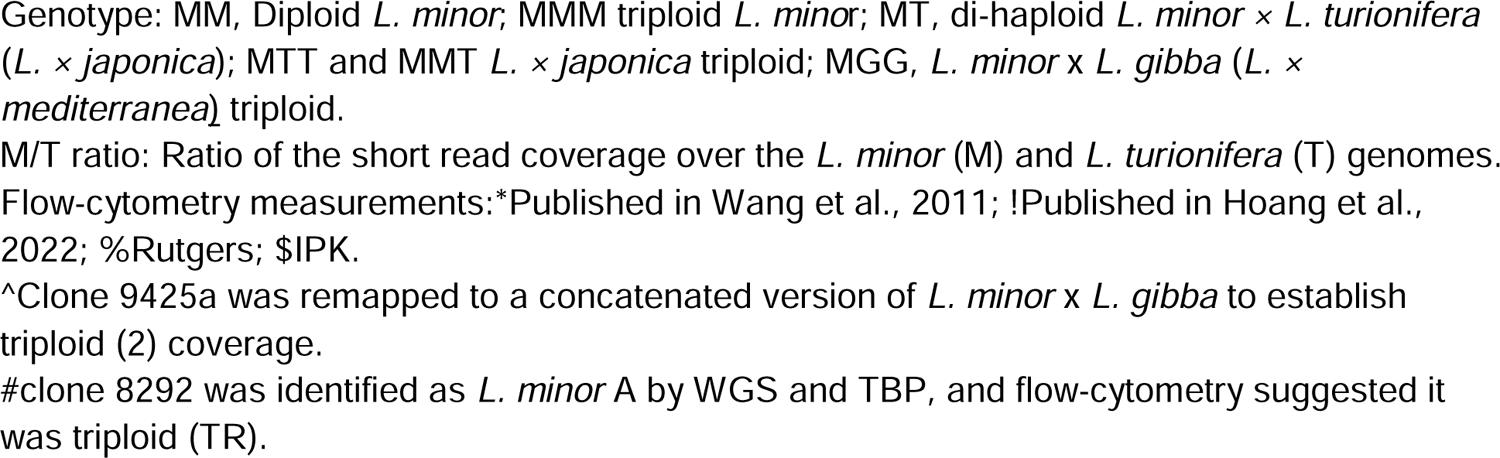
Presumed Lemna minor clones from the Landolt Collection evaluated in the study with the new species designation (predicted species) based on molecular data.

The percent of the genome covered revealed three distinct groups: 1) 55% (32/58) were *L. minor* clones, mapping with high coverage (∼92%) to the *L. minor* genome and low mapping to *L. turionifera* (6–46%); 2) 40% (23/58) were *L. × japonica* (*L. minor × L. turionifera*) hybrids, mapping with high coverage (95%) to both *L. minor* and *L. turionifera*; and 3) two clones (7264, 5635) deviated from this pattern, mapping with lower coverage to both genomes consistent with their low overall mapping rates (Table 1, S1, S2). Clone 8292 with the overall mapping rate of 11%, covered the *L. minor* and *L. turionifera* genome at 91% and 6% respectively, suggesting that it was in fact *L. minor*; however, the low overall mapping rate was due to bacterial contamination (∼88% *Stenotrophomonas*). Since clones 7264 and 5635 had lower overall mapping rates and low (∼7%) coverage of the *L. minor* and *L. turionifera* genomes, we mapped it against other published duckweed genomes and found that both clones mapped at a high overall rate and coverage to *Landoltia punctata* (Baggs *et al*., 2022) (Table 1, S1,2). These results confirm that the presumed *L. minor* clones in the RDSC have been misidentified, revealing a hybrid (9425a) and a different genus (7264, 5635).

Next, we looked at coverage depth across the clones to estimate ploidy since we have shown that *L. × japonica* makes di-haploid and triploid hybrids (Ernst *et al*., 2023), which could explain the differences in genome size estimates. All of the *L. minor* clones mapped with a coverage depth ∼53% to the *L. minor* genome and ∼2% to the *L. turionifera* genome, while in contrast, the *L. × japonica* clones mapped at 33% and 22% respectively. Therefore, all clones had coverage depth close to the target of 60x, except clone 8292 that has a significant amount of bacterial reads. The *L. × japonica* clones further separated based on the ratio of *L. minor* (M) to *L. turionifera* (T) genome coverage depth into ratio of ∼1, ∼2, and ∼0.5, which suggested that we have 7 MT di-haploids, 15 MMT triploids and one MTT triploid (Ernst *et al*., 2023). Therefore, the presumed *L. minor* clones were 55% *L. minor*, 39% *L. × japonica*, 3.4% *Landoltia punctata*, and 1.7% *L. × mediterranea*.

### K-mer genome size and heterozygosity estimate

While we successfully clarified the species identity of the 58 presumed *L. minor* clones, we also sought to estimate their genome sizes. Therefore, we performed k-mer frequency-based genome size estimation using Illumina SR sequencing data. The estimated genome sizes ranged from 318 to 741 Mb (Table 1), spanning a wider range than previous flow cytometry estimates (356–604 Mb) (Wang *et al*., 2011). The median heterozygosity rate was 0.68%, with values ranging from 0.07% to 3.05% (Table 1). The k-mer frequency profiles were grouped into three distinct patterns (Figure 1A-C; Figure S2): 1) single peak; 2) double peak with the first peak taller; and 3) double peak with the second peak taller. K-mer profiles with two peaks typically indicate heterozygosity or polyploidy, whereas a single peak suggests a homozygous diploid genome (Michael and VanBuren, 2015). As expected, clones with heterozygosity below 0.2% consistently exhibited a single k-mer peak, while those with higher heterozygosity displayed a secondary peak or hump (Figure 1A-C; Figure S2). The genomes clustered into three K-mer-predicted average genomes sizes of 330, 514 and 723 Mb that matched with the *L. minor* diploid, *L. × japonica* triploid and *L. × japonica* di-haploid respectively (Figure 1D).

**Figure 1.**
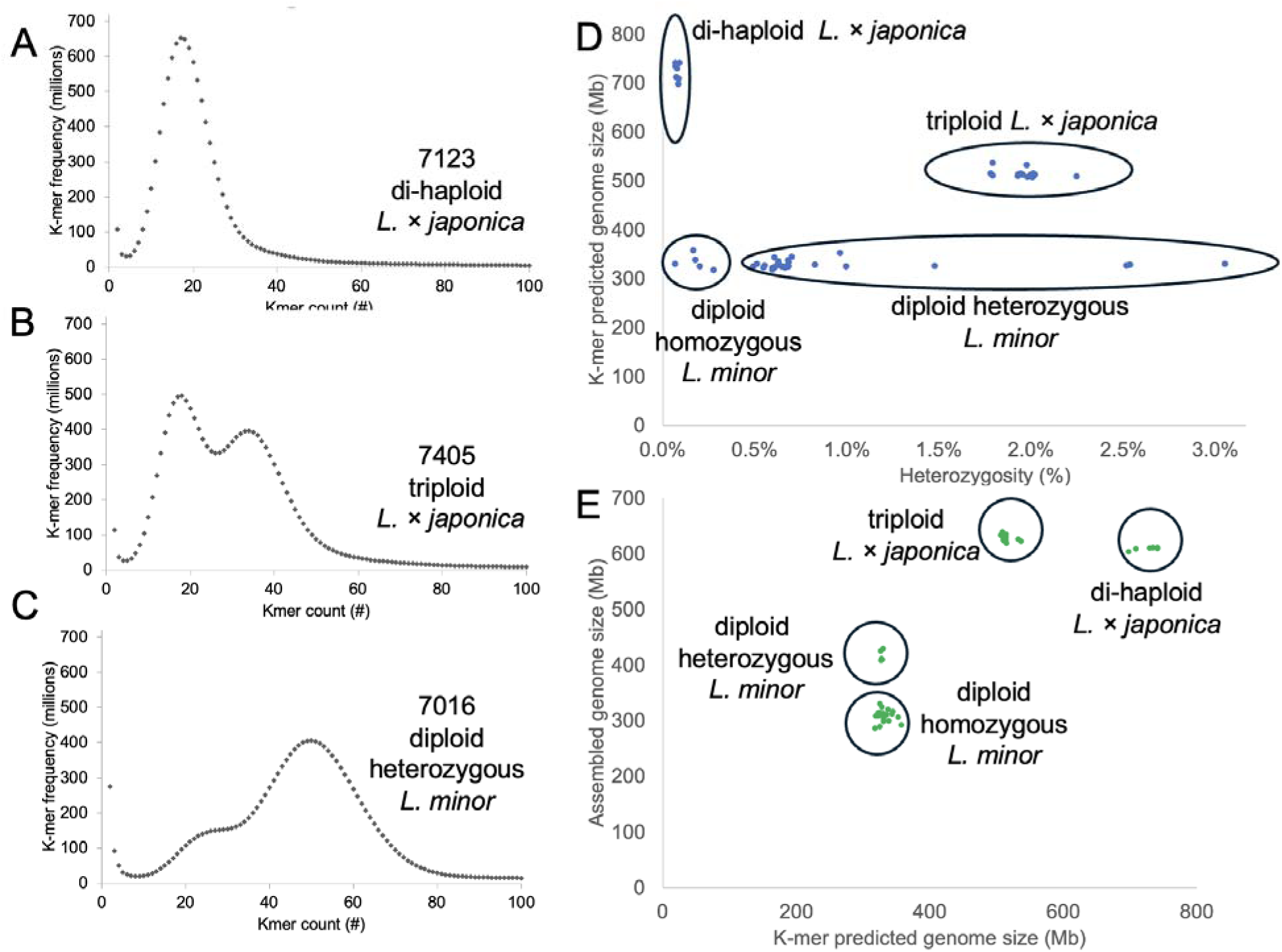
Presumed *L. minor* genome feature estimation using Illumina short reads (SR) and k-mer as well as genome assembly analysis. A-C) K-mer frequency histograms show distinct genome structure patterns for representative clones across different ploidy levels: A) di-haploid (7123) exhibits a single dominant peak, consistent with low heterozygosity; B) triploid (7405) displays a bimodal distribution, indicative of increased heterozygosity; and C) heterozygous diploid (7016) shows a distinct double peak, reflecting higher allelic diversity. D) K-mer-based genome size estimates plotted against heterozygosity (%) reveal four distinct *Lemna* genome classes: homozygous diploids, heterozygous diploids, triploids, and di-haploids. The di-haploid genome exhibited an unusual pattern, with a predicted size twice the base *Lemna* genome size of 350 Mb. This is because *L. minor* and *L. turionifera* are genetically divergent enough that their sequences are recognized as distinct in the k-mer frequency histogram, rather than overlapping as would be expected in a more closely related hybrid. E) Comparison of K-mer-based genome size predictions and assembled genome sizes highlights four genome categories, with triploids exhibiting the largest assembled genome sizes. Species and ploidy designation determined based on the mapping results (Table 1; Table S1,2).

Heterozygosity was almost exclusively associated with the *L. minor* diploid clones, whereas the *L. × japonica* di-haploid exhibited a single peak and low heterozygosity, despite its k-mer-predicted genome size being double that of the *L. minor* and *L. turionifera* base genome (330 Mb) (Figure 1D). Since sequencing coverage confirms that the di-haploid clones contain both *L. minor* and *L. turionifera* genomes, explaining the 2x genome size prediction, their extremely low heterozygosity suggests that the k-mer frequency distribution appears haploid-like. This likely results from the high genetic divergence between the parental genomes, which reduces detectable heterozygosity despite the hybrid nature of the genome. These results in comparison with flow-cytometric genome size determination, chromosome counting and genomic *in situ* hybridization (see below) indicate that relying solely on k-mer analysis may lead to misinterpretation of the data.

### Validating ploidy with orthogonal measures

While we established the ploidy levels of all the clones based on coverage depth, we wanted to see if we could recapitulate these results using a k-mer based approach. Therefore, we leveraged the k-mer based Smudgeplot tool to estimate ploidy levels across the presumed *L. minor* clones (Ranallo-Benavidez *et al*., 2020). The k-mer-based Smudgeplot estimates of ploidy generally agreed with our classifications (Table S1; Figure S3), but there were several important exceptions. For three of the diploid *L. minor* genomes Smudgeplot predicted one as tetraploid (clone 7194) and two as triploid (clones 9533 and 8866). In addition, three of the *L. × japonica* triploids were predicted to be diploids by Smudgeplot (clones 9436, 9541, and 7436). However, Smudgeplot accurately predicted all of the *L. × japonica* di-haploid clones as diploid (Table S1). Smudgeplot is sensitive to sequencing coverage, so it is possible that the discrepancies reflect issues with depth since at least for the triploids the effective coverage is half of that for the others. That means that Smudgeplot by itself was insufficient to definitively assign ploidy levels.

### Flow cytometry genome size estimates

Flow cytometry is a powerful tool for estimating genome size, providing rapid and quantitative measurements of cellular DNA content (Bennett and Leitch, 2005). We performed flow cytometry on a subset of clones to validate our sequencing-based genome size estimates with additional validation from two independent laboratories that maintain these clones separately (Table 1; Table S1). Despite quantitative differences across labs (Table S1), all estimates followed the relative genome size trends observed in the original study (Wang et al., 2011).

Specifically: 1) Predicted *L. minor* diploids had genome sizes of ∼400 Mb, consistent with the 1C haploid base genome size; 2) *L. × japonica* triploids had genome sizes 1.5× larger than diploids (510–650 Mb); 3) *L. × japonica* di-haploids had genome sizes (>400 Mb), somewhat higher than *L. minor* diploids due to the larger genome of *L. turionifera* (Hoang et al., 2022) (Table 1). We also examined flow cytometry results for clones that were potentially misclassified by Smudgeplot, a computational tool for ploidy inference. Clone 9533, initially predicted as a diploid *L. minor* based on k-mer and depth analysis, was identified as triploid by Smudgeplot; a finding that was confirmed by flow cytometry (641 Mb). Clone 7194, classified as a diploid *L. minor* by most methods, was called tetraploid by Smudgeplot but confirmed as diploid by flow cytometry (398 Mb). Two of the three *L. × japonica* triploids that were incorrectly called diploid by Smudgeplot had flow cytometry estimates supporting triploidy (7436: 557 Mb; 9436: 641 Mb). In some cases, flow cytometry results varied between labs, with inconsistencies observed for clones 7018, EL022, 9437, 9440, and 8625 for which the difference is in the scale of a ploidy level. In these cases, most likely different biological samples run under the same number.

Discrepancies at smaller scales can likely be attributed to differences in the used reference species and assumptions made about their genome sizes. If even under the same conditions in one lab different genome sizes for intraspecific accessions appear, these are likely due to changes which accumulated during long periods of clonal propagation. Summarizing these results show flow cytometry as a direct method is the most reliable approach to determine genome size.

### Cytogenetics validate genomics-based results

A robust approach to confirm ploidy status is direct chromosome counting (Hoang *et al*., 2019). We selected a subset of clones for chromosomal visualization to validate our genome size and sequencing-based predictions (Figure 2). As expected, clones 7210 and 7753, representing both homozygous and heterozygous *L. minor* diploids, had 42 chromosomes (2n = 2x = 42).

**Figure 2.**
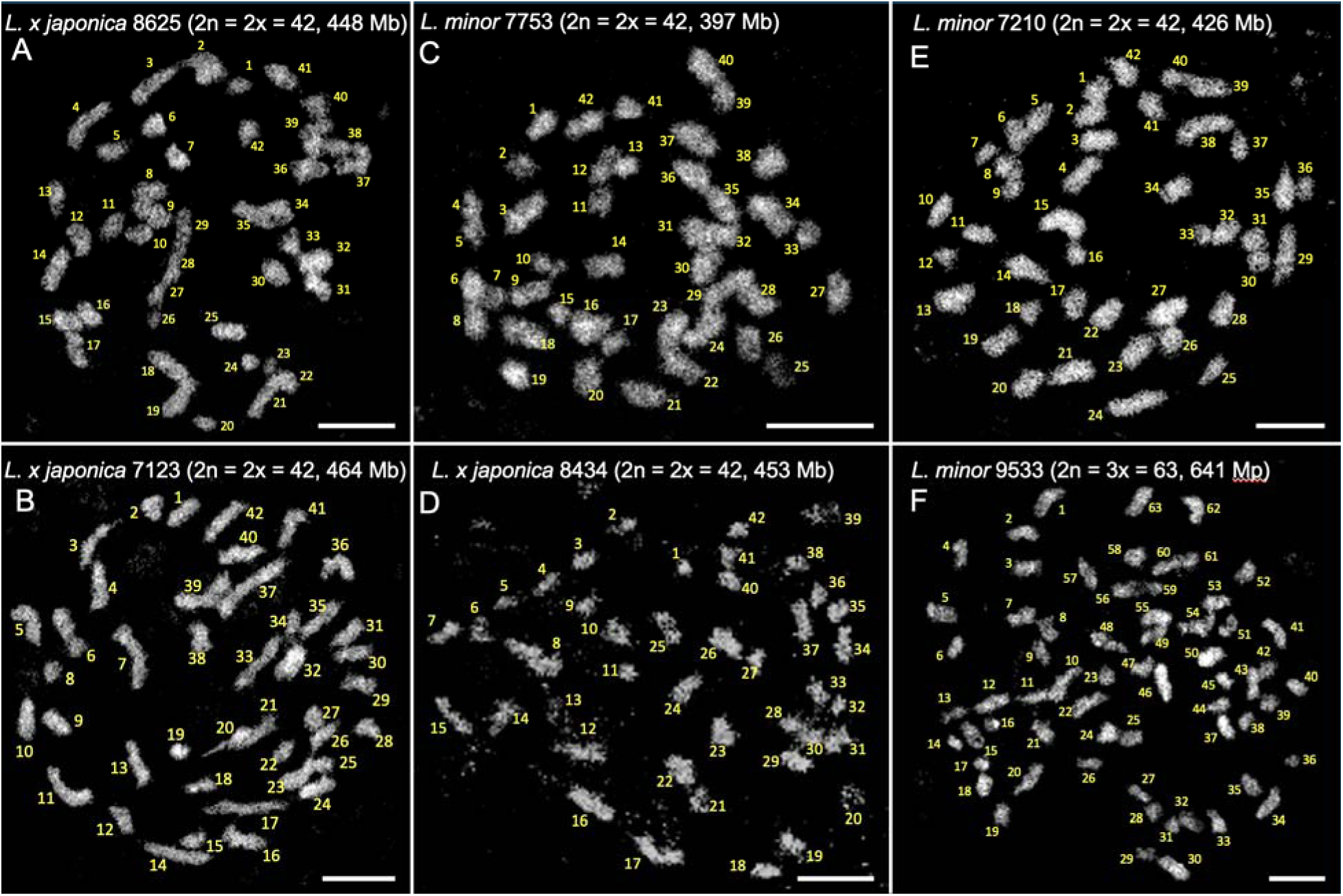
Mitotic chromosome spreads with clones of Lemna. A) *L.× japonica 8625 (2n = 2x = 42, 448 Mbp)* di-haploid; B) *L.× japonica 7123 (2n = 2x = 42, 464 Mbp)* di-haploid; C) *L. minor 7753 (2n = 2x = 42, 397 Mbp)* diploid; D) *L.× japonica 8434 (2n = 2x = 42, 453 Mbp)* di-haploid; E) *L. minor 7210 (2n = 2x = 42, 426 Mbp)* African-clade diploid; F) *L. minor 9533 (2n = 3x = 63, 641 Mbp)* triploid.

However, clone 9533, which appeared diploid based on read depth and k-mer analysis but was classified as triploid by flow cytometry and Smudgeplot, was confirmed to have 63 chromosomes (2n = 3x = 63). Furthermore, clones 8625, 7123, and 8434, all identified as *L. × japonica* di-haploids, consistently exhibited 42 chromosomes (2n = 2x = 42), confirming their di-haploid status.

### Genome assembly to clarify species assignment

*De novo* genome assembly using Illumina SR typically results in modest-sized contigs, yet it effectively captures gene space and provides valuable insights into genome relationships (Michael and VanBuren, 2015). Therefore, we *de novo* assembled the Illumina short reads and found the genomes ranged in quality with N50 lengths spanning from 3 to 21 Kb, which is consistent with assemblies from short reads (Table 1; Table S1). The genomes grouped into two size classes at 300-400 Mb and 600 Mb that matched with *L. minor* diploid and *L. × japonica* di-haploid/triploid respectively (Figure 1E). Among the diploid *L. minor* clones, five individuals (9533, 8211, 7650, 8866, and 7753) had relatively larger assembled genome sizes (400 Mb) and were also predicted to have the highest heterozygosity (1.5–3.1%). The larger assembled genome sizes in heterozygous diploids can be attributed to the incomplete collapse of haplotypes, a common issue in SR assemblies. When heterozygosity is high, haplotypes contain sufficient variation to be assembled as separate contigs, leading to inflated genome size estimates. This pattern is further supported by a strong negative correlation (r = −0.8) between N50 contig length and heterozygosity (Figure S4), indicating that higher heterozygosity results in shorter contigs and larger total assembly sizes due to partial haplotype duplication. The *L. × japonica* di-haploid/triploid assemblies were approximately twice the size of the diploid *L. minor* genomes, consistent with separate assembly of both subgenomes. This interpretation is reinforced by mapping these assemblies back to the *L. × japonica* 8627 reference genome (Ernst *et al*., 2023), which revealed a clear separation of subgenomic sequences (Figure S5).

The genome assembly estimated that the base *L. minor* genome size was ∼350 Mb in contrast to the value estimated by flow cytometry (∼400 Mb), which could reflect the fact that SR assembly does not generally assemble repeat sequences like centromere and transposable elements well. The larger variation observed previously by flow cytometry (Wang *et al*., 2011) was most likely due to *L. × japonica* triploids.

### Reference-free super-pangenome

A super-pangenome extends the traditional pangenome concept by incorporating multiple species within a genus or across related taxa, rather than being confined to a single species (Raza *et al*., 2023). We leveraged the highest-quality 38 SR *de novo* genome assemblies to construct a reference-free super-pangenome (Figure 3); we also leveraged published chromosome-scale genomes to place the super-pangenome in the context of *L. × japonica*, *L. turionifera* as well as other published duckweed genera (Ernst *et al*., 2023; Todd P. Michael *et al*., 2020; Michael *et al*., 2017; Abramson *et al*., 2021; Baggs *et al*., 2022) (Figure S6). Using PanKmer (Aylward *et al*., 2023), we built a shared k-mer matrix to assess genome-wide relationships across the diploid *L. minor* and *L. × japonica* di-haploid and triploid clones (Figure 3). The reference-free super-pangenome matrix revealed three primary clusters, with a clear outlier grouping of *Landoltia punctata* (clones 5635 and 7264). Additionally, *L. × mediterranea* (clone 9425a) branched separately with the known diploid *L. minor* clone Lmin5500 (Van Hoeck *et al*., 2015), supporting its distinct hybrid status. A particularly useful validation came from clones DWC123 and 7283, which were descendants of the same clone independently maintained by two stock centers for over at least two decades. Their close clustering in the super-pangenome matrix confirms the reliability of our approach in capturing genomic relationships.

**Figure 3.**
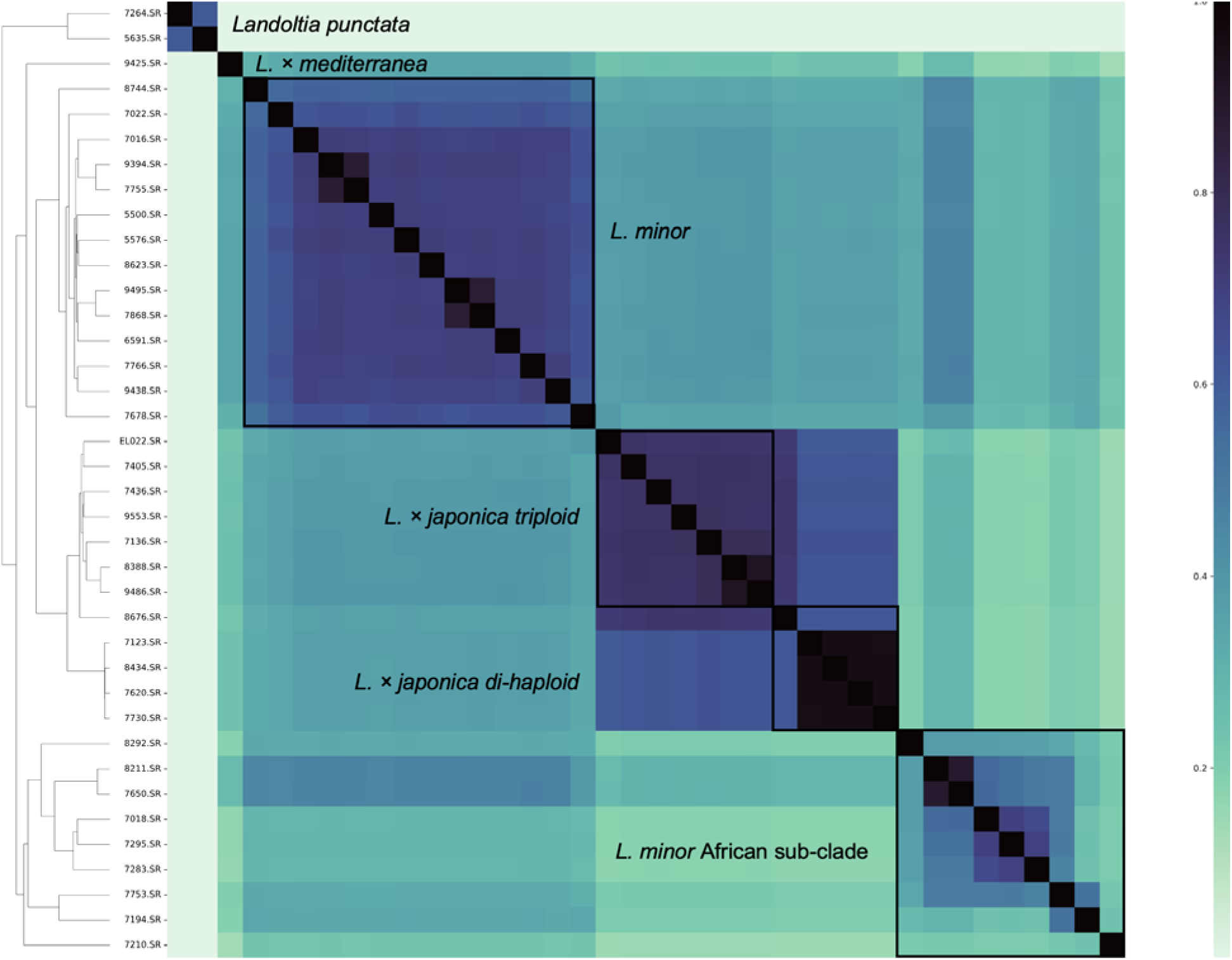
Reference-free super-pangenome based on Illumina short read (SR) arranged presumed *L. minor* genomes into species and ploidy classes based on k-mer distance. A jaccard super-pangenome matrix was generated with Illumina short read (SR) based genome assemblies using PanKmer (K=31). Darker color corresponds to less k-mer distance, while lighter color corresponds to more k-mer distance. *Landoltia punctata*, *L.× mediterranea*, *L. mino* diploids, *L.× japonica* triploids, *L.× japonica* di-haploids, and *L. minor* African-clade are boxed.

The three main clusters in the super-pangenome included two on either side of the matrix that contained the diploid *L. minor* clones, while the third cluster in the middle contained the *L. × japonica* di-haploids and triploids (Figure 3). The top left *L. minor* diploid cluster included the published Lmin5500 clone (Van Hoeck *et al*., 2015), which suggested that the bottom right *L. minor* diploid cluster might represent a distinct sub-class. Moreover, the top left *L. minor* diploid cluster shared more k-mers with the *L. × japonica* di-haploid/triploid cluster than the other diploid cluster; this supported that the diploid clusters may represent different subclasses as well as suggested that these *L. minor* diploids were the sub-class that contributed to the *L. × japonica* hybrids. When we included all published duckweed chromosome-scale genomes in the super-pangenome, the *L. × japonica* di-haploid/triploid cluster further supported the hybrid nature with *L. turionifera* and suggested that the *L. × mediterranea* hybrid 9445a may have shared k-mers with clone 7210 from the *L. minor* African-clade (Figure S6). These findings demonstrate that super-pangenome matrices provide a powerful genome-wide approach for rapidly assessing relationships between clonal populations within a species and even across genera. The super-pangenome approach enabled a reference-free comparison of genome structure and hybridization patterns, which enhanced our ability to differentiate subclasses within *L. minor*, track hybrid origins, and explore genome evolution at a broader taxonomic scale.

### Tubulin intron-Based Polymorphism (TBP) validation

*Tubulin intron-based polymorphism (TBP)* calling is another approach that has been utilized to clarify the relationship between *Lemna* species as well as to identify hybrids derived from *L. minor* and *L. turionifera* called *L.× japonica (Braglia, Lauria, et al., 2021; Volkova et al., 2023; Braglia, Breviario, et al., 2021)*. All of the presumed *L. minor* clones (except the *Landoltia* and *L.× mediterranea* clones) were validated using the TBP marker; both cluster as well as PCA analyses were consistent with three groupings similar to the super-pangenome matrix (Figure 4; Table S1). The top left cluster from the super-pangenome matrix overlapped with the true *L. minor* diploid clones similar to Lmin5500 (*L. minor* 5500-like; Figure 4). The center super-pangenome cluster overlapped with the *L.× japonica* cluster as predicted. However, the TBP analysis suggested clone 7868, which matched with diploid *L. minor* based on sequencing, was *L.× japonica*. The most parsimonious explanation is that this also reflected a clone name representing different biological material across labs. The third TBP cluster that was distant compared to the other two clusters (*L. minor* African-clade), overlapped with the lower right cluster of *L. minor* diploids of the super-pangenome, which we also found more distant in the k-mer matrix. All these clones come from Africa and some Middle Eastern countries such as Iran and Turkey. Therefore, we have termed this subclass the “African-clade.” Considering the diversity of this cluster in the super-pangenome matrix, this cluster could represent different species or hybrids, but distinct from the recently identified *L.× mediterranea* clones (Braglia *et al*., 2024; Romano *et al*., 2024).

**Figure 4.**
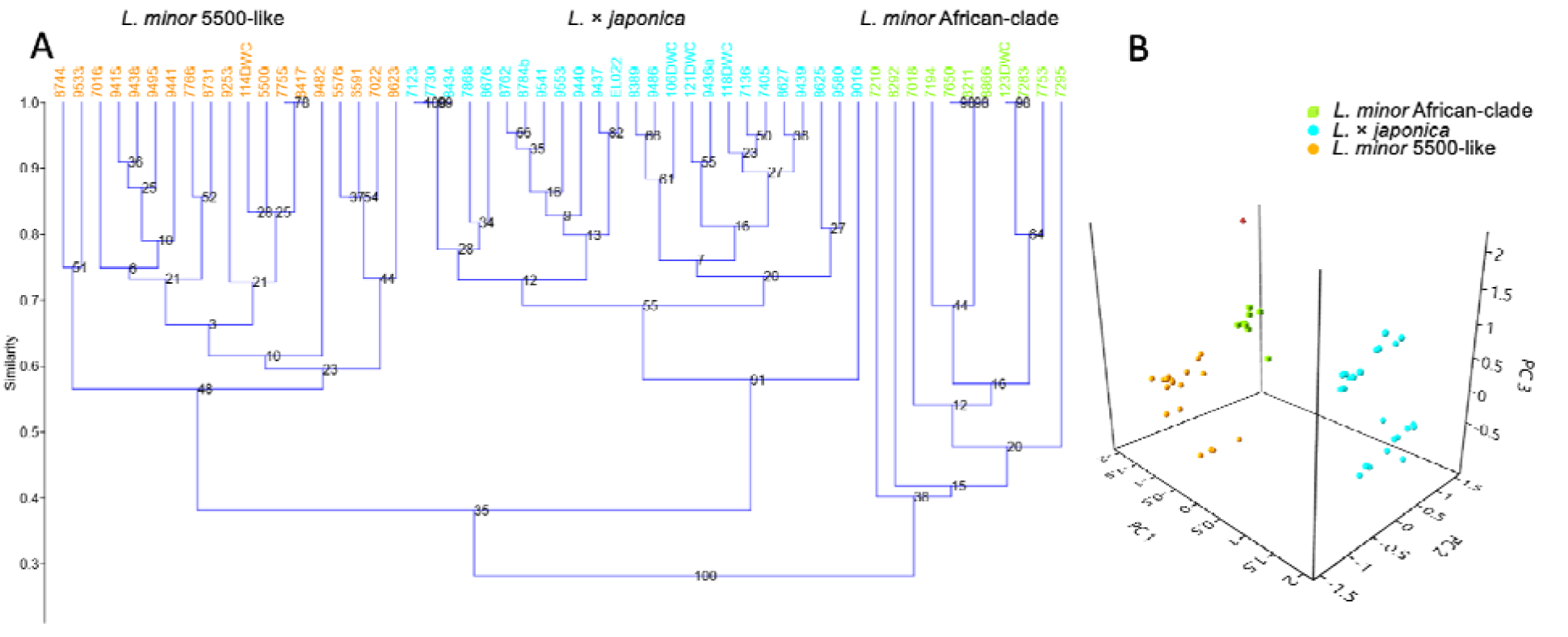
TBP analysis confirms two *Lemna* species, *L. minor* and *L. x japonica*, and suggests a further *L. minor* African-clade. A) Jaccard UPGMA tree based on the β-tubulin intron size variation. B) PCA plot of the relationship between *L. minor* clones using *Landoltia punctata* as outgroup (red symbol).

### Confirmation of the hybrids by Genomic In Situ Hybridization (GISH)

We performed GISH to confirm the ploidy status and genome components of *L. × japonica* triploids and di-haploids. Using species-specific probes designed from *L. minor* (M) and *L. turionifera* (T) genomes, we visualized the sub-genomes at the chromosome level (Figure 5). For the *L. × japonica* di-haploid clone 8434 (Ljap8434), GISH confirmed a diploid chromosome number (Figure 2D; 5A), reinforcing its classification. Clone 9533, originally identified as diploid based on k-mer analysis and read depth, was later predicted as triploid by Smudgeplot and flow cytometry. Our GISH analysis validated the triploid classification (2n = 3x = 63; 641 Mb/1C) by demonstrating that all chromosomes were predominantly labeled by *L. minor* DNA (Figure 5B). This confirms that 9533 is a *L. minor* autotriploid (see also Figure 4), not a hybrid, explaining why it initially clustered with diploid clones. Notably, 9533 and 8292 are the only confirmed *L. minor* triploids in our dataset. Additionally, GISH confirmed that clone 8627 is a *L. × japonica* triploid (2n = 3x = 63) (Ernst et al., 2023), further supporting our ploidy classifications. These results reinforce the importance of combining sequencing-based and cytogenetic methods to accurately classify ploidy and genome composition, particularly in complex hybrid species.

**Figure 5.**
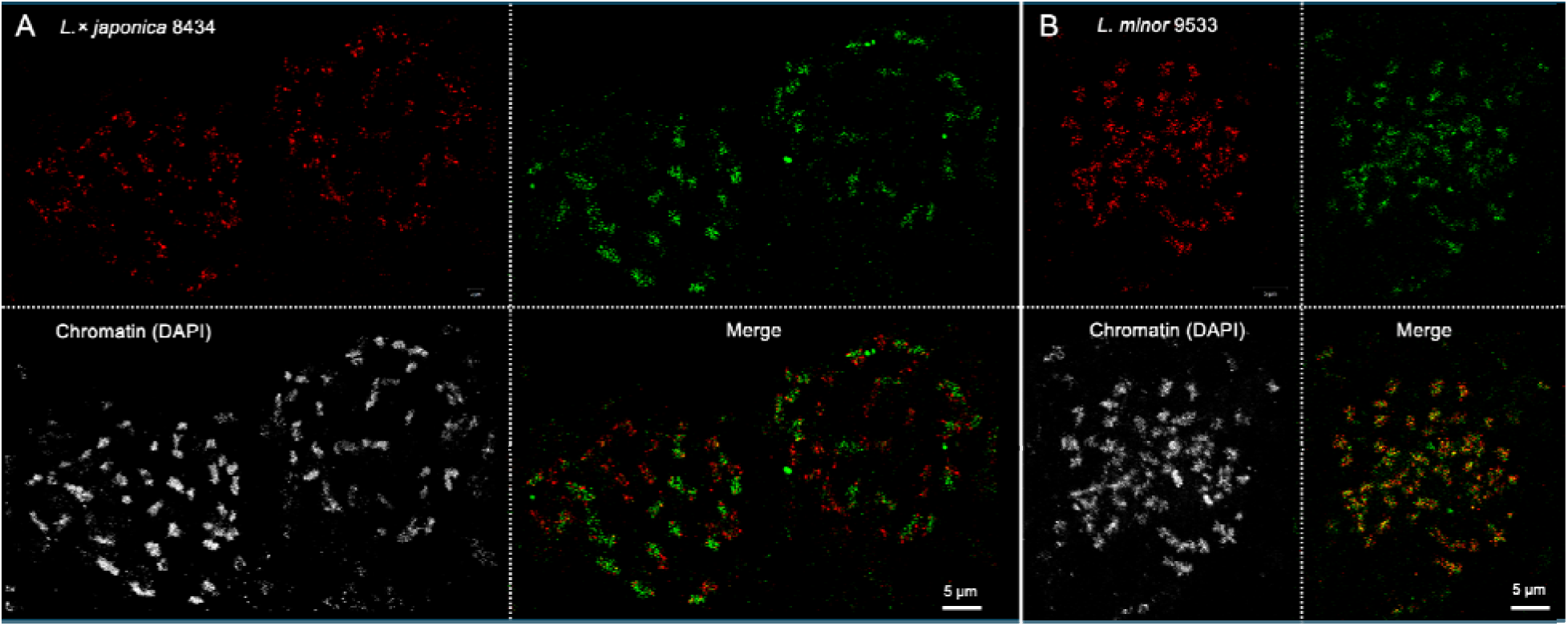
GISH on somatic metaphase chromosomes of the di-haploid *L.× japonica* 8434 (A) and the autotriploid *L. minor* 9533 (B). GISH was performed with genomic DNA of *L. minor* (red) and *L. turionifera* (green).

### Summary of presumed L. minor clone identity

Integrating results from multiple approaches and independent laboratories, we reclassified the 58 presumed *L. minor* clones based on their species identity and ploidy level (Table 1; Table S1). We found that 55% were *L. minor* clones that separated into two sub-classes (5500-like and 7210-like), and that clones 9533 and 8292 were the only *L. minor* triploids in the collection. However, 45% of the collection were mis-called. Most of them were reassigned to *L. × japonica* di-haploids (12%) and triploids (28%) (Table 1; Table S1). We only found one clone (7425a) in the collection (1.7%) that was a hybrid between *L. minor* and *L. gibba* (*L. × mediterranea*).

Finally, we identified two clones (3.4%) that were from the monospecific genus *Landoltia* (*Landoltia punctata*), suggesting that confusion in assigning clone identity across the *Lemna* complex may extend to other genera. These findings underscore the importance of integrating multiple genomic and cytogenetic tools for accurate species and ploidy classification, as misidentifications remain common in collections of morphologically similar duckweed species.

### Polyploids provide growth advantage in some environments but not in others

Polyploidy is hypothesized to enhance vigor and environmental adaptability, whereas diploids are often specialized for narrow ecological niches (Freeling, 2017). In *Lemna*, both interspecific and intraspecific hybrids may benefit from increased growth rate and environmental flexibility due to heterosis and genomic buffering (Chen and Birchler, 2013). We measured 24 clones, which included diploid and polyploid *Lemna* species, regarding their growth parameters: relative growth rate (RGR), doubling time (DT), relative yield (RY), and mean area (MA) (Ziegler *et al*., 2015). Both *L. × japonica* di-haploid and triploid lines exhibited superior growth performance across most traits. Di-haploids outperformed diploids in DT, RY, and RGR, but showed the smallest effect on MA (Table 2; Table S3). The triploid *L. minor* clone (9533) displayed the largest mean area (MA, P < 0.05) among all classes, suggesting a size advantage associated with triploidy. In contrast, diploid *L. minor* clones were the most variable and exhibited lower overall vigor, reinforcing the hypothesis that polyploidy contributes to greater robustness and growth potential.

**Table 2.** Growth measurements across the *Lemna* inter- and intraspecific hybrids and diploids.

|  | RGR (day<sup>-1</sup>) | DT (days) | RY (wk<sup>-1</sup>) | MA (cm<sup>2</sup>) |
| --- | --- | --- | --- | --- |
| Di-haploid (*L. x japonica*) N = 4 | 0.48 ± 0.01 | 1.46 ± 0.04 | 30.74 ± 2.54 | 1.14 ± 0.04 |
| Diploid African-clade (*L. minor* A) N = 4 | 0.37 ± 0.02 | 2.08 ± 0.09 | 16.6 ± 1.96 | 1.26 ± 0.08 |
| Diploid (*L. minor*) N = 7 | 0.41 ± 0.01 | 1.71 ± 0.06 | 19.72 ± 1.55 | 0.96 ± 0.07 |
| Triploid (*L. x japonica*) N = 8 | 0.46 ± 0.02 | 1.53 ± 0.06 | 26.9 ± 3.44 | 1.52 ± 0.06 |
| Triploid (*L. minor*) N = 1 | 0.42 ± 0.02 | 1.71 ± 0.07 | 22.24 ± 2.72 | 1.78 ± 0.05 |

Polyploids are hypothesized to exhibit greater vigor and adaptability across a wider range of environmental conditions (Wendel, 2015). To test this, we measured relative growth rate (RGR), relative yield (RY), and doubling time (DT) for one representative clone from each class (*L. minor* diploid, *L. × japonica* di-haploid, and *L. × japonica* triploid) under five different light and temperature conditions at 7 and 14 days (Figure 6; Tables S4, S5). Growth was assessed at two time points since earlier growth rates tend to be faster and more linear with less plant crowding and higher nutrient availability. Significant differences (P > 0.1) were observed for both condition and clone effects on RGR and RY, while DT showed no significant associations (Figure 6; Figure S7; Tables S4, S5). Under control conditions, *L. × japonica* di-haploids and triploids grew 24% and 35% faster than the *L. minor* diploid. A larger growth advantage emerged under low-light conditions, where the hybrids exhibited 81% (di-haploid) and 51% (triploid) higher growth rates compared to the diploid. Importantly, under higher temperature or high light stresses, the di-haploids and triploids both showed less fitness compared to the diploids. These findings suggest that polyploidy may confer a growth advantage under specific environmental conditions, particularly low-light stress, but may be less tolerant to more extreme abiotic stressors such as high temperature or light.

**Figure 6.**
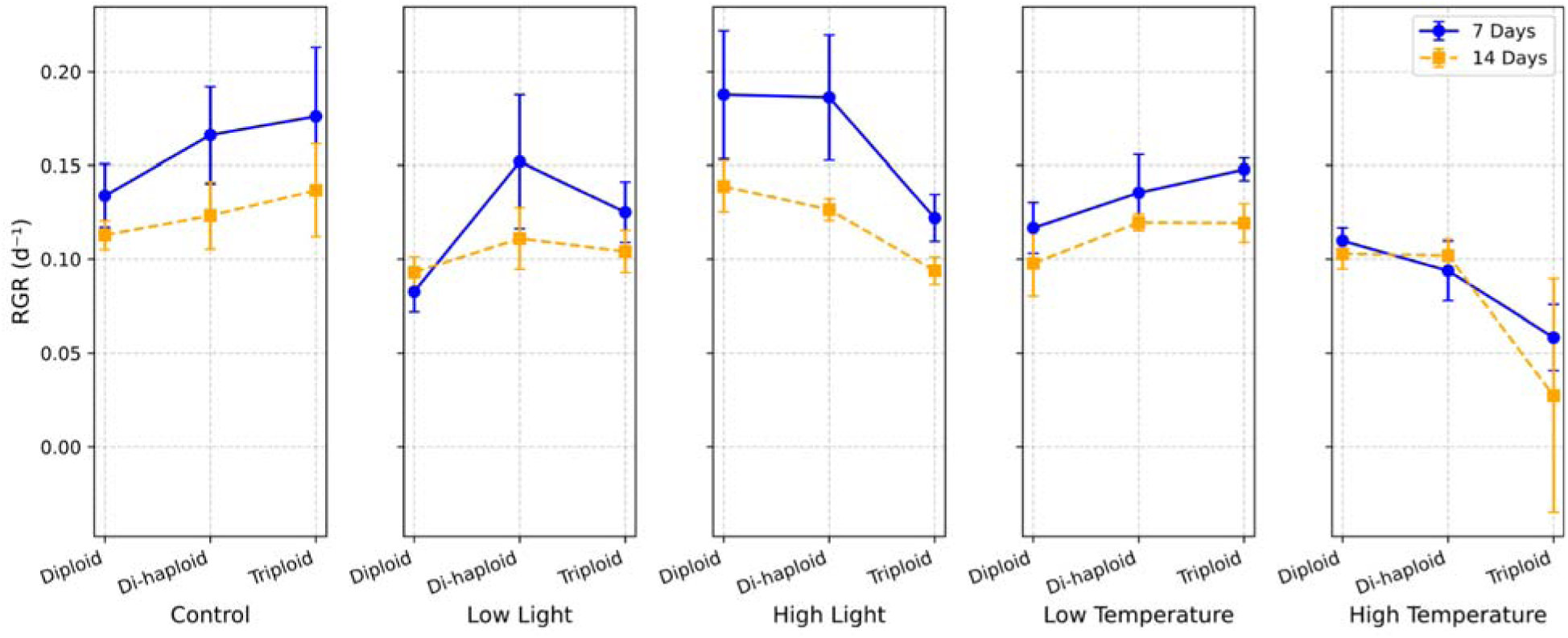
Relative growth rate (RGR) of diploid, di-haploid and triploid *Lemna* clones across variable light and temperature conditions. RGR was measured at 7 days (d, blue) and 14 d (orange) for a diploid *L. minor* (7766), *L. x japonica* di-haploid (8676) and *L. x japonica* triploid (9440) under control (150 μmol m^−2^ s^−1^; 25L°C), low light (50 μmol m^−2^ s^−1^; 25L°C), high light (220 μmol m^−2^ s^−1^; 25L°C), low temperature (150 μmol m^−2^ s^−1^; 15L°C) and high temperature (150 μmol m^−2^ s^−1^; 35L°C).

## Discussion

*Lemna minor*, known as common duckweed, is widely regarded as the most abundant and broadly distributed duckweed species on Earth (Landolt, 1986). However, the widespread ability of *L. minor* to colonize diverse nutrient- and water-rich habitats suggests that this morphologically similar yet genetically diverse group may actually represent a “*Lemna minor* complex,” consisting of closely related species and hybrids (Braglia *et al*., 2021; Braglia *et al*., 2024; Schmid *et al*., 2024; Braglia *et al*., 2021; Volkova *et al*., 2023). In this work, we leverage a multi-pronged approach, comprising WGS, k-mer analysis, flow-cytometry, TBP molecular marker, chromosome-counting and GISH to characterize the species and ploidies of the previously presumed 58 *L. minor* clones from the Landolt Duckweed Collection. Our results re-classified these *L. minor* clones into: 1) two classes of diploid *L. minor* clones that includes a grouping we propose to call the African-clade (*L. minor A*); 2) di-haploid and triploid *L.× japonica* (*L. minor x L. turionifera*) hybrids; 3) one triploid *L. minor × L. gibba* (*L. × mediterranea*); and 4) two clones from the *Landoltia* (7264, 5635) genera. In addition, we found two *L. minor* triploid clones (8292, 9533), one of them was validated by GISH and TBP (9533) (Figs. 4 and 5) and the other (8292) by TBP. Moreover, now that we could clearly group the clones by species and ploidy, we were able to show that the polyploids display growth advantages under specific conditions but not under the abiotic stressors of high light and temperature. These results set the stage for proper assignment of future clones added to the collection as well as enable future research on how polyploidy impacts duckweed growth and expansion.

Currently the Landolt Duckweed Collection is hosted at two locations: the Rutgers Duckweed Stock Cooperative (RDSC; http://www.ruduckweed.org/), and the National Research Council, CNR, Milan, Italy (https://biomemory.cnr.it/collections/CNR-IBBA-MIDW). Therefore, we assessed genome size and the TBP molecular marker in clones maintained across different labs. Our analysis revealed several discrepancies between clones from different laboratories, a phenomenon also observed in the model plant *Arabidopsis thaliana*. Notably, even the widely used reference line Columbia-0 (Col-0) has been shown to accumulate distinct genomic variations when maintained in separate labs (1001 Genomes Consortium, 2016). For instance, we found that flow-cytometry results differed by lab for five clones (7018, EL022, 9437, 9440, and 8635), most likely reflecting either experimental mislabeling or the same accession number representing different biological samples (Table S1). Only one clone (7868) differed between the WGS and TBP, which most likely also reflected a mislabeling between labs. However, the two clones (DWC123 [UTCC270], and 7283) that were separated by at least two decades in two different collections, but originated from the Landolt Duckweed Collection, were exactly the same (Figure 3). These results highlight that in order to get reliable results multiple approaches, including direct ones in addition to probabilistic ones, are essential, in particular when looking at species complexes like the *Lemna minor* complex.

A puzzling phenomenon remains: triploids and interspecific hybrids, both of which arise only through sexual reproduction, occur relatively frequently in populations that predominantly propagate asexually (Acosta *et al*., 2021; Mateo-Elizalde *et al*., 2023). Polyploidy predominantly arises either by fusion of unreduced gametes or after spontaneous chromosome doubling, resulting in allopolyploids if di-haploid hybrids were involved (Wang *et al*., 2019). Triploids appear after fusion of a reduced with an unreduced gamete and can be maintained only by asexual propagation, because irregular meiotic chromosome segregation does not yield viable gametes. The apparent loss of genes involved in the small RNA-mediated triploid block pathway as well as genes that prevent unreduced gametes seem to be responsible for the frequent occurrence of triploids (Ernst *et al*., 2023). Moreover, the predominantly asexual propagation habit of duckweeds would be consistent with the avid viability of triploids arising in duckweeds. We found that 40% (23) of the previously identified *Lemna minor* strains (based predominantly on morphological basis) we screened in the RDSC collection were in fact a mixture of *L.× japonica* with 70% (16) of those being triploid and 30% (7) di-haploids (Table 1, S1). A similar situation was found for presumed wild *L. minor* clones collected across Switzerland where over 50% were identified as *L.× japonica* triploids by WGS. Their low genetic variability indicated that triploids may emerge, clonally multiply and exploit specific environments (Schmid *et al*., 2024). Similarly, among wild presumed *L. minor* clones from Eastern Europe, molecular markers were used to show that *L.× japonica* was prominent (Volkova *et al*., 2023). Apparently, sexual hybrids, including triploids, are more common than previously thought in nature, suggesting that *L.× japonica* may confer some level of advantage in particularly adapted habitats as observed in our growth experiments (Figure 7; Figure S7; Tables S3-5).

Duckweeds exhibit a wide range of genetic diversity and reproductive strategies that influence their adaptability to different environments. Some species, such as *Spirodela polyrhiza*, maintain large population sizes but low sequence variation (Xu *et al*., 2019), suggesting that even minor genetic advantages can enhance clonal propagation and facilitate the colonization of new micro-environments. Similarly, wild populations of presumed *L. minor* have been found to harbor 4 to 20 distinct genotypes per site, reflecting extensive genetic diversity that may aid in local adaptation (Bog *et al*., 2022). Supporting this, *L. minor* exhibits twice as much genetic variation in fitness per site compared to other species, consistent with strong purifying selection acting to counteract environmental fluctuations (Jewell and Bell, 2023). While hybrids (di-haploids and triploids) were not explicitly resolved in these previous studies, their presence could contribute to local adaptation and population dynamics. Our findings align with this framework, as hybrid genotypes outperformed diploids under specific conditions but exhibited reduced growth under abiotic stressors of high temperatures and light (Figure 6, S7; Table 2, S3-5). A similar pattern has been reported for *L. minor* and *L. x japonica* clones (Usui and Angert, 2024), reinforcing the idea that hybridization may offer a competitive advantage in stable environments but limit resilience under environmental stress. Although the growth advantage of triploids appears modest (yet significant), even a 5% increase in relative growth rate (RGR) would allow a particular clone to rapidly outcompete and dominate a population over time.

In conclusion, our comprehensive characterization of presumed *L. minor* clones from the Landolt Duckweed Collection underscores that these clones form the “*Lemna minor* complex” that includes *L. minor, L. turionifera* and *L. gibba,* autotriploids, as well as di-haploid and reciprocal triploid interspecific hybrids (Braglia *et al*., 2024; Ernst *et al*., 2023), rather than a uniform species, and are far more genetically diverse than previously recognized. Similarly, a “*Lemna aequinoctialis* complex” has recently been described (Stepanenko *et al*., 2025), suggesting that the *Lemna* genus may exploit its environment through various hybrid strategies. The discovered hybrid and polyploid clones, apparently arose rather frequently among the rare sexual progenies and under ‘standard’ conditions outperform diploids. Thus, sexuality plays a more pivotal role in duckweed evolution than once assumed. These findings extend our understanding of how polyploidy and hybridization may influence ecological success and genetic diversity in duckweeds, providing a base for future work on their evolutionary dynamics, taxonomy, and biotechnological applications.

## Methods

### Duckweed clones

*Lemna minor* clones were received from the Rutgers Duckweed Stock Collective (RDSC; http://www.ruduckweed.org/). Clones were checked for sterile conditions and if they were not sterile they were re-sterilized per the RDSC protocol. 58 clones were grown under 12 h light and 12 h dark at 22°C. The collection location and all associated information are reported in *Table S1*.

### DNA extraction

Clones were collected, patted dry to remove excess water and flash frozen in liquid nitrogen. Frozen tissue was ground in a mortar and pestle under liquid nitrogen. HMW DNA was isolated using a modified BOMB protocol (https://bomb.bio/protocols/). DNA quality was assessed on a bioanalyzer and HMW status was confirmed on an agarose gel.

### Genome sequencing

Illumina 2×150 bp paired end reads were generated for genome size estimates, short-read genome assembly and polishing genome long read assemblies. Libraries were prepared from HMW DNA using NEBnext (NEB, Beverly, MA) and sequenced on the Illumina NovaSeq (San Diego, CA). Resulting raw sequence was only trimmed for adaptors, resulting >60x coverage of the diploid *L. minor* genome (350 Mb).

### Variation estimation by read mapping

Illumina paired end reads were mapped to the *L. minor* diploid reference genome (Lmin5500) (Van Hoeck *et al*., 2015), as well as the updated *L. minor* 9252 and *L. turionifera* 9434 (Ernst *et al*., 2023) using minimap2 (Li, 2018). Clone 9425a was also mapped to a concatenated version of *L. minor* 9252 and *L. gibba* 7742a. Mapping rates were summarized in Table S1,2. Resulting sorted bam files were analyzed for mapping statistics using samtools (Danecek *et al*., 2021) and variants were called using samtools mpileup (Li *et al*., 2009) and vcftools (Danecek *et al*., 2011).

### Genome size estimation by K-mer frequency

K-mer (k=31) frequency was estimated with Illumina paired end reads (2×150 bp) using Jellyfish (v2.3.0) (Marçais and Kingsford, 2011) and analyzed using in house scripts and GenomeScope and GenomeScope2 (Vurture *et al*., 2017; Ranallo-Benavidez *et al*., 2020). Genome size, heterozygosity and repeat content were first estimated with GenomeScope (Vurture *et al*., 2017), which reports haploid genome size (Table 1, S1). Ploidy was estimated using GenomeScope2 and smudgeplot (Ranallo-Benavidez *et al*., 2020) (Table 1, S1).

### Short read genome assembly

Illumina paired end reads were assembled with Spades (v3.14.0) with standard settings (Bankevich *et al*., 2012). Resulting contigs were mapped to the *L. minor* reference genome (Lmin5500) to estimate completeness and statistics were determined (Table 1, S1).

### Reference-free Pangenome (PanKmer)

The short read-based genome assemblies were used to construct a pangenome using PanKmer (Aylward *et al*., 2023). Briefly, a 31-mer index was constructed using PanKmer (Aylward et al. 2023) with default parameters with the short read Illumina assemblies. Jaccard and ANI similarities for all assemblies in both indexes were calculated and visualized using PanKmer by setting the clustermap metric parameter to “--metric jaccard” and “--metric ani”, respectively.

### Flow cytometry (FC) genome size estimates

#### Rutgers

The methods used to prepare the nuclei suspensions were based on the protocol for estimation of nuclear DNA content in plants using flow cytometry as described (Dolezel *et al*., 2007). Sixteen clones of *L. minor* and four additional plant species of known ploidy, *Spirodela polyrhiza* (158 Mb), *Arabidopsis thaliana* (134 Mb), *Brachypodium distachyon* (272 Mb), and *Fragaria x ananassa* (800 Mb), were analyzed. These species, along with two of the *L. minor* strains known to be diploid, were used as standards to aid in ploidy estimation of the remaining fourteen *L. minor* varieties. For each strain and standard, 120 mg of fresh tissue was harvested and placed in a petri dish with 4 mL of Galbraith’s buffer, a solution consisting of 45mM MgCl_2_, 20mM MOPS, 30 mM sodium citrate, and 0.1% (vol/vol) Triton X-100. The tissue was gently chopped into small pieces with a disposable razor blade to release nuclei. After the homogenate was filtered through a 40 μm cell strainer, the filtrate was evenly distributed into four microcentrifuge tubes. Propidium iodide was used to stain the DNA; 50μL each of 1mg/mL PI and 1 mg/mL RNase A were added to each of three tubes, the fourth left unstained to act as the control. The samples were incubated on ice for about 15 min and gently shaken periodically before analysis through the flow cytometer. The flow cytometer was set to collect data on 10,000 events, with the minimum core size and flow rate due to the small size of the nuclei. The selected threshold was 80,000.

#### IPK

The genome size of 26 Lemna clones was determined following the procedure according to Dolezel et al. (2007) using the CyStain PI Absolute P reagent kit (Sysmex-Partec) according to the manufacturer’s instructions. Fresh frond tissue was co-chopped with fresh leaf tissue of one of the internal reference standards, either *Raphanus sativus* ‘Voran’ (Gatersleben Gene bank accession: RA 34; 1.11 pg/2C) or *Lycopersicon esculentum* ‘Stupicke Rane’ (Gatersleben Gene bank accession: LYC 418; 1.96 pg/2C). The nuclei suspensions were measured either on a CyFlow Space flow cytometer (Sysmex-Partec) or on a BD Influx cell sorter (BD Biosciences). Each clone was measured at least four times on two different days.

### Tubulin intron-based polymorphism (TBP) analysis

The TBP amplification protocol and data analysis were performed, from 30 ng of total genomic DNA, according to the procedure as described (Braglia *et al*., 2023). The polymorphism in length of both intron regions (I and II) of the L-tubulin genes were scrutinized, and each DNA sample was independently analyzed twice.

### Growth rate and frond area estimation

Clones for relative growth rate (RGR; day^-1^), doubling time (DT; day), relative weekly yield (RY; week^-1^), and mean area (MA; cm^2^)estimation were acclimatized by pre-cultivation for four weeks under axenic conditions in a climate cabinet (PK-520, poly klima GmbH, Freising, Germany) under continuous white light (100 µmol m-2 s-1 PAR) and constant temperature of 25°C ± 1°C. The cultivation medium (N medium) was changed weekly to avoid nutrient deficiency (Appenroth *et al*., 1996). For the main experiment, a colony consisting of three to four fronds was placed in a plastic cup containing 40 ml of N medium. After seven days, the number of fronds was recorded again and the relative growth rate was calculated as RGR = (ln x_t7_ – ln x_t0_) / (t7 – t0) (Ziegler *et al*., 2015). The frond area was measured as the mean value of three single fronds per clone in six replicates using a digital microscope (VFX 7000, Keyence, Neu-Isenburg, Germany).

### Duckweed growth rate estimation under differential conditions

Lemna minor clones were obtained from RDSC (Rutgers Duckweed Stock Collection) at Rutgers the State University of New Jersey, New Brunswick, NJ, USA. The clones were maintained on 0.5X Schenk-Hildebrand medium with 0.8% w/v agar and 0.3% sucrose with the addition of 100 mg/L cefotaxime. The clones were kept at 25°C under illumination of 150 μmol m^−2^s^−1^ light (16 h light/8 h dark). Five frond colonies of L. minor clones from stock cultures were grown for two weeks in a solution containing 0.5X Schenk-Hildebrand medium and 0.3% sucrose. For growth measurement, ∼100 mg fresh fronds of each clone were then transferred to a 177 ml glass jar containing 50 ml of fresh 0.5X Schenk-Hildebrand medium and 0.3% sucrose solution. The experiment was carried out in 3 replicates with samples collected at days 7 and 14. Tissues of L. minor clones were harvested and dried overnight at 55°C in 1.5 ml Eppendorf tubes and the growth rates measured (Relative Growth Rate (RGR; day−1), Doubling time (DT; day), Relative yields (RY; week−1) according to procedures as described (Ziegler et al., 2015). For all experiments, the L. minor clones were maintained at 25°C under illumination of 150 μmol m^−2^s^−1^ light (16 h light/8 h dark). A two-way analysis of variance (ANOVA) was performed using the Statsmodels library in Python to estimate the effects of Condition, Clone, and Time Point on RGR, DT, and RY, after confirming data normality (Shapiro-Wilk test) and equal variance across groups (Levene’s test).

### Chromosome counting

The fronds were grown in liquid nutrient medium under 16 h white light of 100 µmol m-2 s-1 at 24°C (Appenroth *et al*., 1996). The mitotic chromosome spreading was carried out according to (Hoang and Schubert, 2017). Fronds were treated in 2 mM 8-hydroxylquinoline, fixed in fresh 3:1 absolute ethanol: acetic acid, softened in PC enzyme mixture [1% pectinase and 1% cellulase in Na-citrate buffer, pH 4.6], macerated and squashed in 45% acetic acid. After freezing in liquid nitrogen, chromosome spreads were treated with pepsin [50 µg/ml in 0.01 N HCl], post-fixed in 4% formaldehyde in 2x SSC [300 mM Na-citrate, 30 mM NaCl, pH 7.0], rinsed twice in 2x SSC, 5 min each, dehydrated in an ethanol series (70, 90 and 96%, 2 min each) air-dried. Chromosome counting was performed within different slices of 3D-SIM image stacks (Hoang *et al*., 2022).

### DNA isolation

For each sample, 0.3 g of fresh and healthy fronds were harvested and cleaned in distilled water, put into a 2 ml Eppendorf tube with two metal balls, frozen in liquid nitrogen, and ground by a ball mill mixer (Retsch MM400). The genomic DNA of the studied species was isolated using the DNeasy Plant Mini Kit (cat. nos. 69104-Qiagen). DNA was eluted by 200 µl buffer AE and quality checked by electrophoresis. Genomic DNA was sonicated before labelling.

### Probe preparation

Sonicated genomic DNA (1 µg) was labeled with Cy3-dUTP (GE Healthcare Life Science) or Alexa Fluor 488-5-dUTP (Life Technologies) by nick-translation, then precipitated in ethanol (Mandáková and Lysak, 2008) with sonicated unlabeled DNA of the other presumed parental species as carrier DNA in excess. Probe pellets from 10 µL nick translation product for GISH probes were dissolved in 100 µL hybridization buffer [50% (v/v) formamide, 20% (w/v) dextran sulfate in 2× SSC, pH 7] at 37°C for at least 1 h. The ready-to-use probes were stored at −20°C.

### GISH (Genomic in situ hybridization)

Probes were pre-denatured at 95°C for 5 min and chilled on ice for 10 min before adding 20 µL probe per slide. Two-rounds of GISH with alternatively labeled genomic probes of the presumed parental species were performed to investigate the distribution of the corresponding probe signals on the chromosome complement of the tested clones as described (Hoang *et al*., 2022).

### Super-resolution microscopy

To analyze the ultrastructure and spatial arrangement of signals and chromatin at a lateral resolution of ∼120 nm (super-resolution, achieved with a 488 nm laser), spatial structured illumination microscopy (3D-SIM) was applied using a Plan-Apochromat 63x/1.4 oil objective of an Elyra PS.1 microscope system and the software ZENblack (Carl Zeiss GmbH) (Weisshart *et al*., 2016).

## Data availability

Raw Illumina short read sequencing data were deposited under BioProject PRJNA602906.

## Author contributions

T.P.M., E.L. and B.W.A. conceived the project. I.P. and E.L. maintain the *L. minor* collection. K.C. and M.L. grew, sterilized, and extracted DNA from *L. minor* clones. S.P. processed short read sequencing data, estimated genome size, assembled short reads, mapped short reads, estimated variation and constructed pangenome. P.T.N.H., V.S., and I.S. provided chromosome counts and GISH, and J.F. and M.W. estimated genome size using flow cytometry. K.S.S., K.A., M.B., B.P. and E.L. conducted growth rate experiments. T.P.M. wrote the manuscript and the co-authors edited. L.B. and L.M. validated species designation using the Tubulin intron-Based Polymorphism (TPB) molecular markers.

## Acknowledgements and funding

This work was funded in part by the Tang Genomics Fund to T.P.M. Work on duckweed in the Lam lab and support for the Rutgers Duckweed Stock Cooperative are supported by a grant from the Gordon and Betty Moore Foundation (AWD00009516) via a subaward from the University of Toronto. In addition, this work was supported by a Hatch project (12116) and a Multi-State Capacity project (NJ12710) from the New Jersey Agricultural Experiment Station at Rutgers University (B.P., E.L.). This work was partially supported by the Italian National Research Council within the Agritech National Research Center and received funding from the European Union Next Generation EU [grant ID Piano Nazionale Di Ripresa e Resilienza (PNRR), Missione 4 Componente 2, Investimento 1.4—Project CN00000022].

## Supplemental Tables

Table S1. Full table of *L. minor* clones.

Table S2. Mapping coverage across the presumed *L. minor* clones against *L. minor* and *L. turionifera*.

Table S3. *L. minor* and *L. × japonica* di-haploid and triploid growth rates.

Table S4. *Lemna* growth characteristics under different conditions.

Table S5. ANOVA of the growth measures RGR, DT and RY across environmental conditions.

## Supplemental Figures

**Figure S1.**
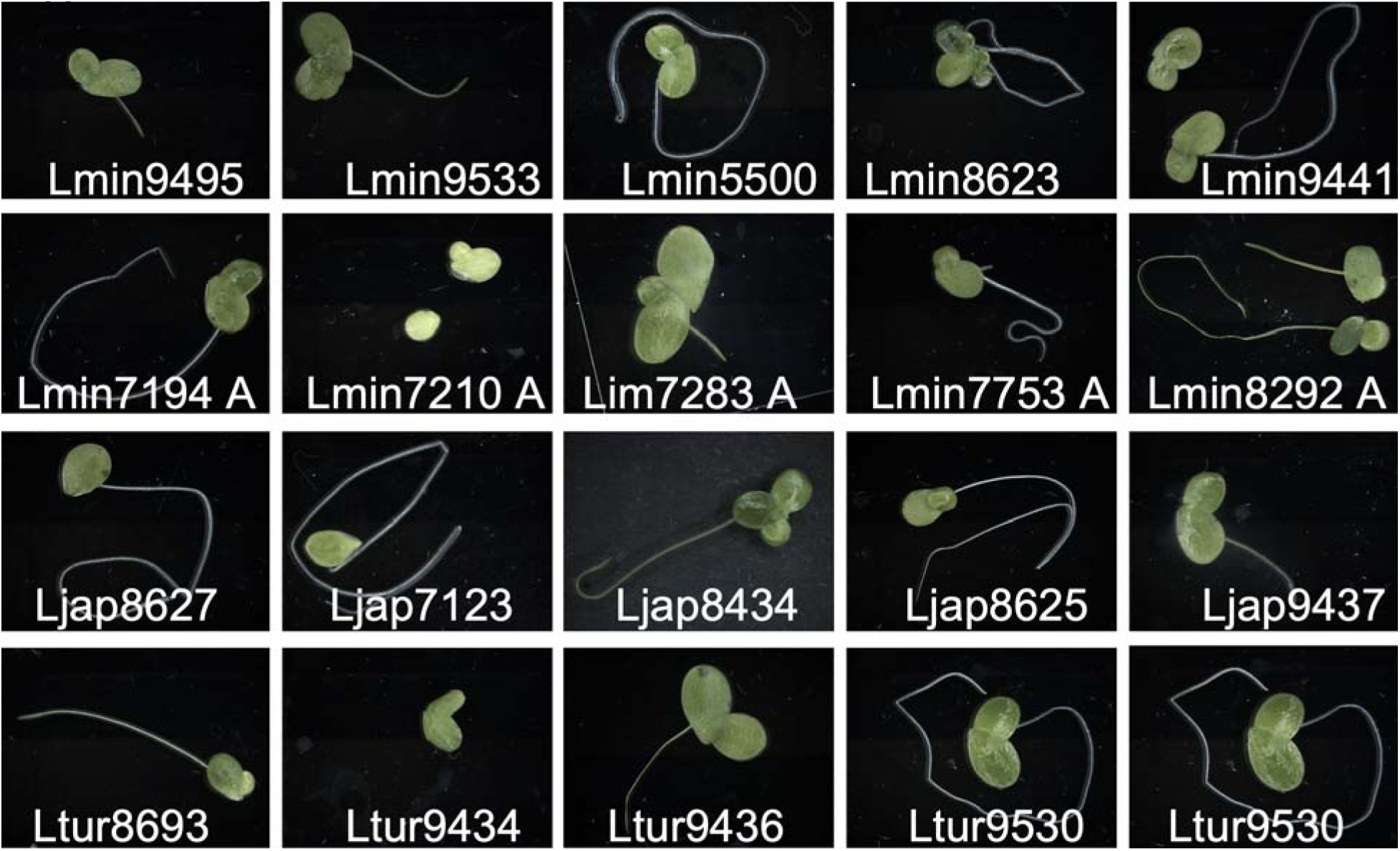
Presumed *L. minor* clones from the Landolt Duckweed Collection stored at the Rutgers Duckweed Stock Collective (RDSC) are difficult to tell apart. Putative *L. minor* (Lmin) and *L. turionifera* (Ltur) clones from the Landolt Duckweed Collection, maintained at the Rutgers Duckweed Stock Cooperative (RDSC), exhibit morphological similarity, making them difficult to distinguish visually. The *L. turionifera* clones were included for contrast but were not tested in this study. This highlights the challenges of species identification within the “*Lemna minor* complex,” emphasizing the need for genomic and molecular approaches to accurately resolve species identity and hybrid classification. The species designation is from Tables 1, S1. LminXXXX A clones are part of the African-clade.

**Figure S2.**
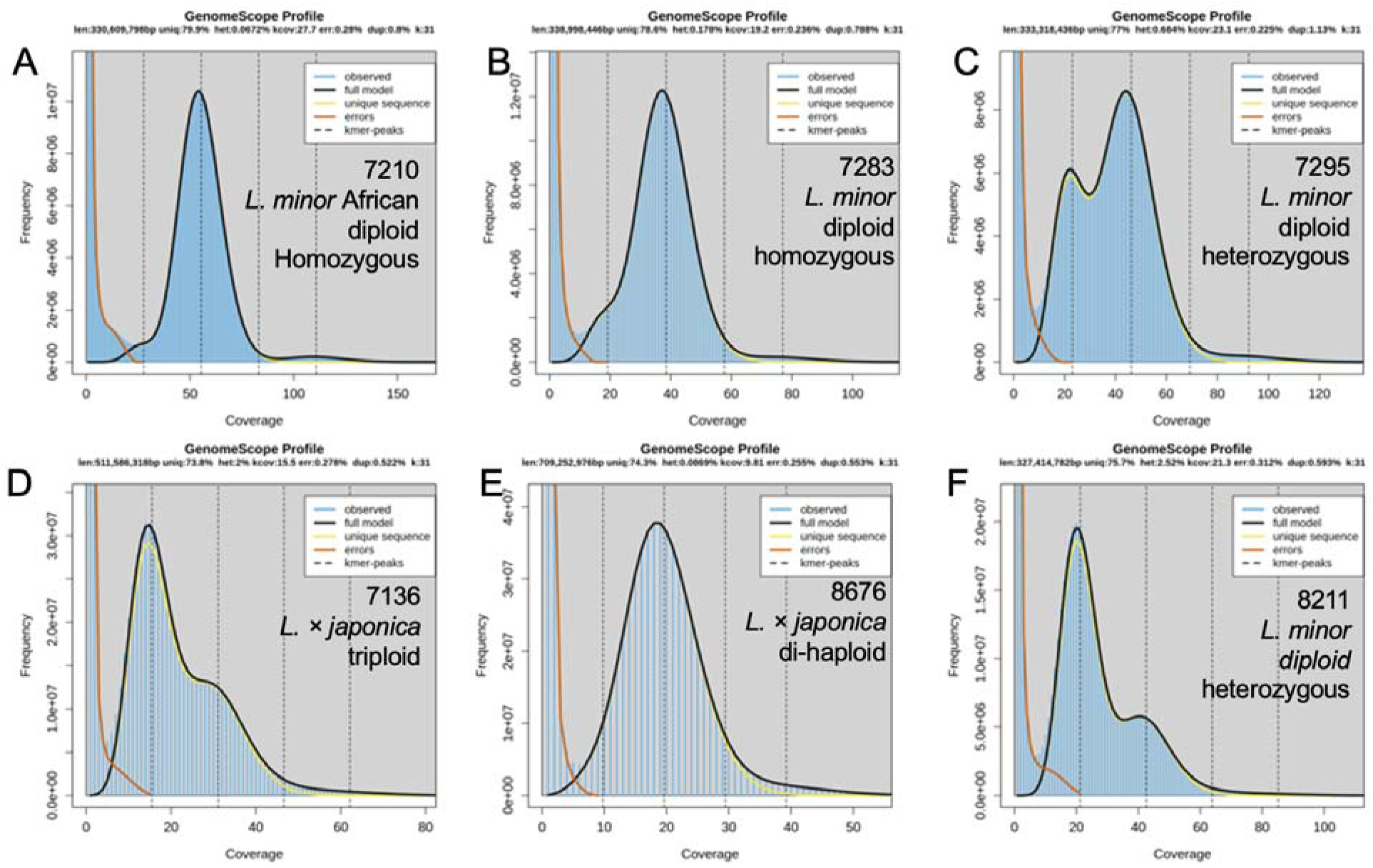
*L. minor* k-mer (k=31) frequency histograms from GenomeScope. K-mer frequency histograms (K=31) generated using GenomeScope suggest genome size, heterozygosity, and repeat composition across the presumed *L. minor* clones. Genome size is inferred from the total k-mer distribution. Heterozygosity is estimated based on the presence of distinct peaks, where higher secondary peak prominence indicates increased heterozygosity. Repeat structure is reflected in the proportion of highly repetitive k-mers contributing to overall genome complexity. Panels represent histograms for six presumed *L. minor* clones: A) 7210; B) 7283; C) 7295; D) 7136; E) 8676; F) 8211. Species and ploidy were determined based on mapping results (Table 1; Table S1,2).

**Figure S3.**
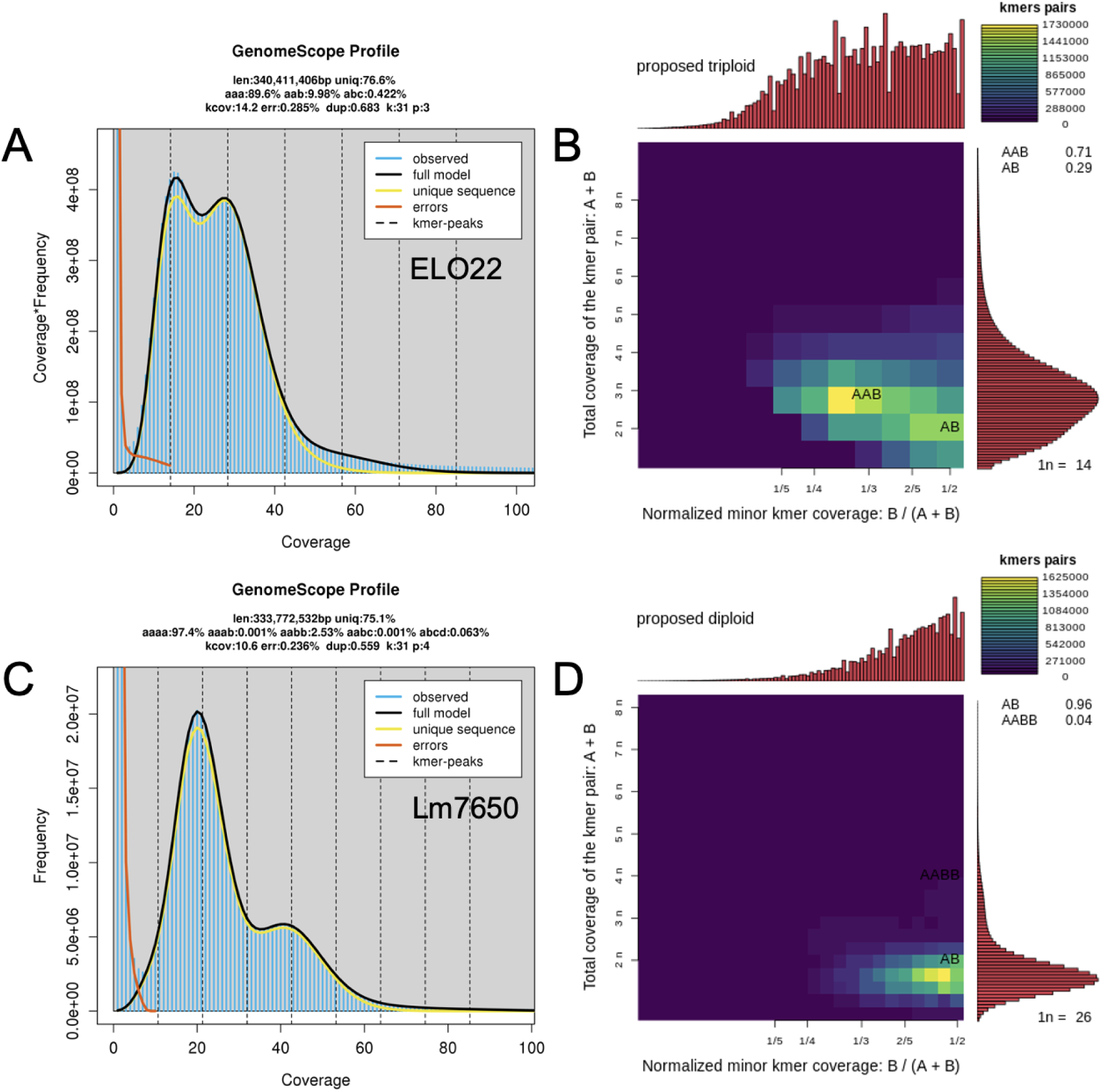
Examples of ploidy estimated using GenomeScope2 and Smudgeplot. A) Clone ELO22, which is estimated to be a triploid, is confirmed using GenomeScope2 using triploid settings, and B) is predicted to be triploid by Smudgeplot. C) Clone 7650 was predicted by read mapping to be *L. minor* and the k-mer plot suggested high heterozygosity and the first peak being higher than the second peak. D) Smudgeplot predicted clone 7650 as a diploid.

**Figure S4.**
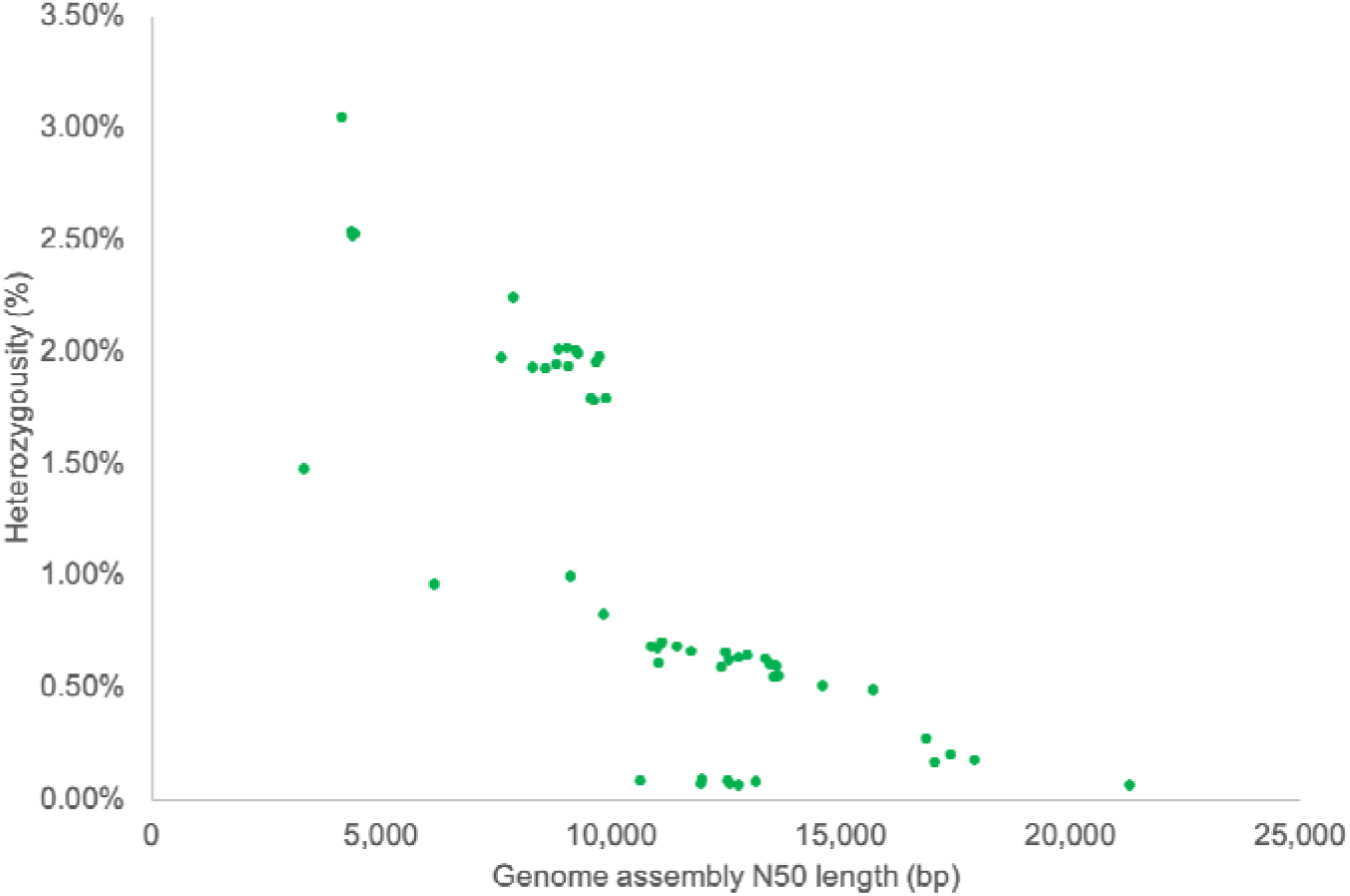
Correlation between *de novo* genome assembly N50 length and k-mer predicted heterozygosity. This figure illustrates the inverse relationship between *de novo* genome assembly N50 length (bp) and k-mer predicted heterozygosity. As heterozygosity increases, the N50 length decreases, reflecting the limitations of short-read assemblies in resolving highly heterozygous regions. Higher heterozygosity leads to a more fragmented assembly as divergent haplotypes create complex assembly graphs, causing contig splitting. Lower heterozygosity allows for longer, more contiguous assemblies, as there are fewer sequence variants disrupting scaffold construction. These results highlight the impact of genome heterozygosity on assembly contiguity, emphasizing the challenges of reconstructing structurally complex genomes using short-read sequencing.

**Figure S5.**
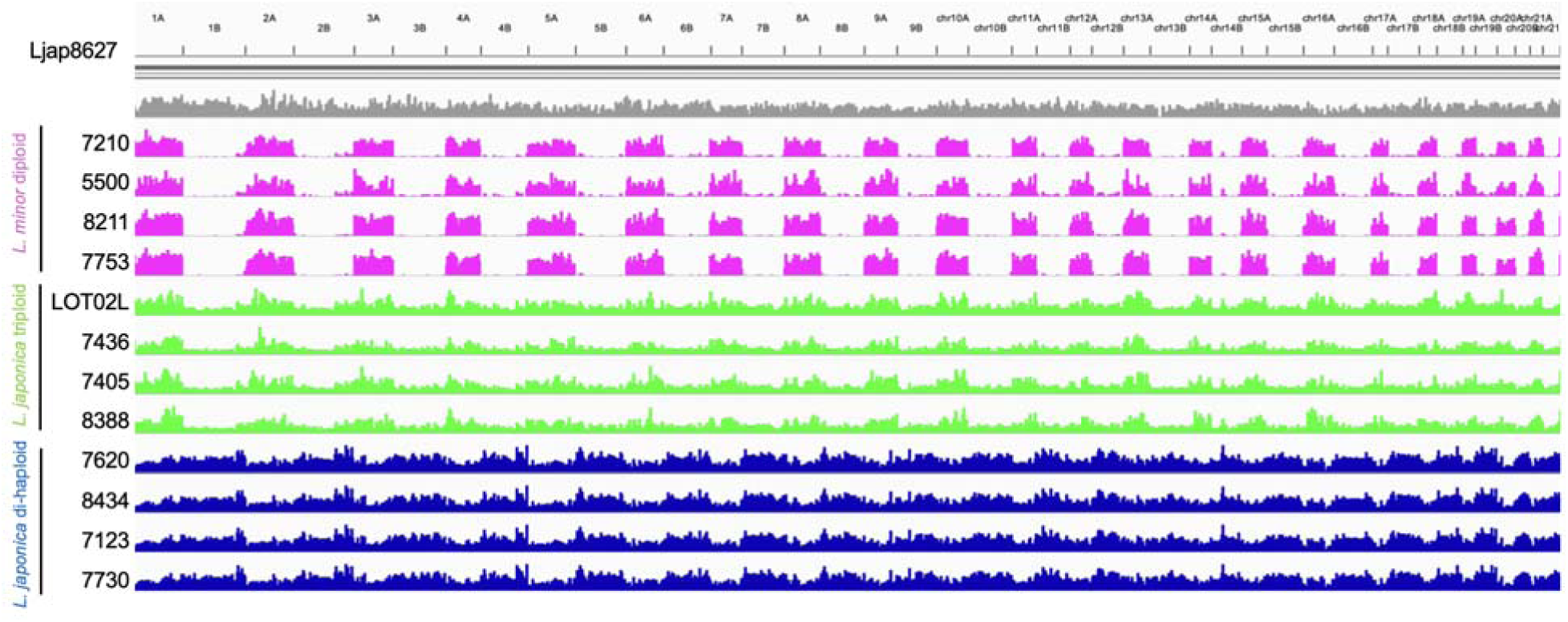
Mapping of short read (SR) genome assemblies to the *L. × japonica* Ljap8627 chromosome-scale reference genome. SR genome assemblies from *Lemna* diploids, triploids, and di-haploids were mapped to the chromosome-scale *L. × japonica* Ljap8627 reference genome (Ernst *et al*., 2023) to assess their alignment patterns across sub-genomes. Top track (gray): Coverage profile of the Ljap8627 reference genome, which represents a hybrid between *L. minor* and *L. turionifera*. Pink tracks: *L. minor* diploids (7210, 5500, 8211, 7753), which map strongly to the *L. minor* sub-genome, showing distinct coverage peaks corresponding to the haploid genome structure. Green tracks: *L. × japonica* triploids (LOT02L, 7436, 7405, 8388), which exhibit a more continuous coverage profile, reflecting contributions from both *L. minor* and *L. turionifera* sub-genomes. Blue tracks: *L. × japonica* di-haploids (7620, 8434, 7123, 7730), which display even coverage across both sub-genomes, consistent with a balanced hybrid genome composition. The mapping patterns confirm the hybrid nature of *L. × japonica* triploids and di-haploids, which contain sequences from both *L. minor* and *L. turionifera*. The *L. minor* diploids map primarily to their sub-genome, showing a distinct, punctuated coverage pattern. The higher coverage uniformity in triploids and di-haploids suggests that they inherit structural and genomic contributions from both parental species, reinforcing their hybrid classification.

**Figure S6.**
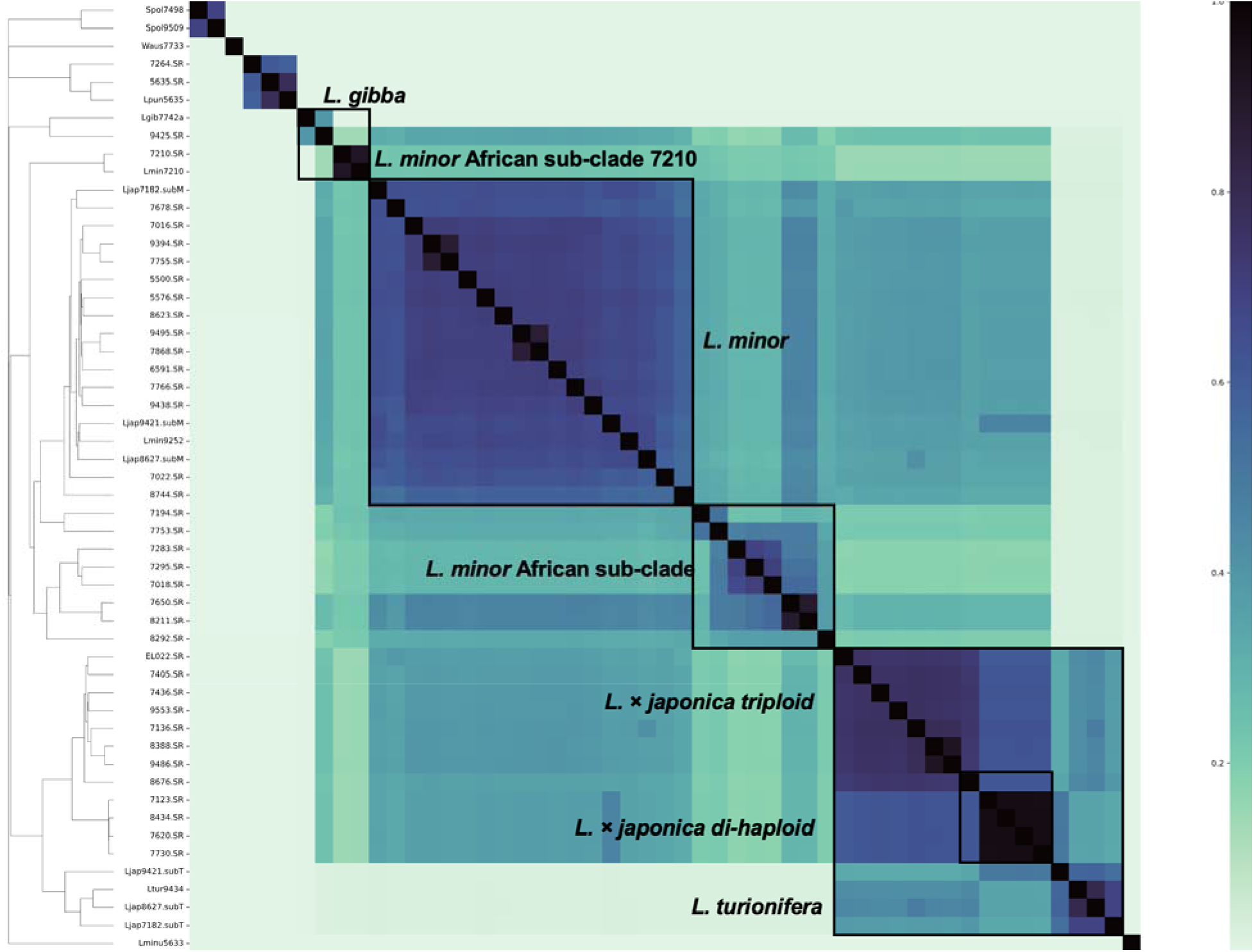
Reference-free super-pangenome arranged presumed *L. minor* genomes into species and ploidy classes based on k-mer distance. A Jaccard super-pangenome matrix was constructed using Illumina short-read (SR) genome assemblies alongside published chromosome-scale duckweed genomes, leveraging PanKmer (K=31) to assess genomic similarity based on k-mer distance. Darker colors indicate greater similarity (lower k-mer distance), while lighter colors reflect higher genomic divergence. The inclusion of chromosome-scale genomes shifts cluster positions in the matrix compared to the SR-only analysis, as sub-genomes separate (*L. minor* subM, *L. turionifera* subT) and additional reference genomes (*L. turionifera* 9434, *L. gibba* 7742a) refine species classifications (Figure 3). The hybrid clone 9425a (*L. × mediterranea*) clusters with *L. gibba* 7742a, one of its sub-genomes, and also shifts *L. minor* African-clade clone 7210 into closer proximity, suggesting shared k-mers from its L. minor sub-genome. The chromosome-scale *L. turionifera* reference genome further clarifies the relationship between *L. × japonica* triploids and di-haploids, confirming their hybrid origins. These findings highlight the power of reference-free super-pangenomes in distinguishing species and ploidy levels based on k-mer similarity, providing new insights into sub-genome contributions in hybrid duckweed lineages. Previously published chromosome-scale genomes (Ernst *et al*., 2023; Todd P. Michael *et al*., 2020; Michael *et al*., 2017; Abramson *et al*., 2021; Baggs *et al*., 2022).

**Figure S7.**
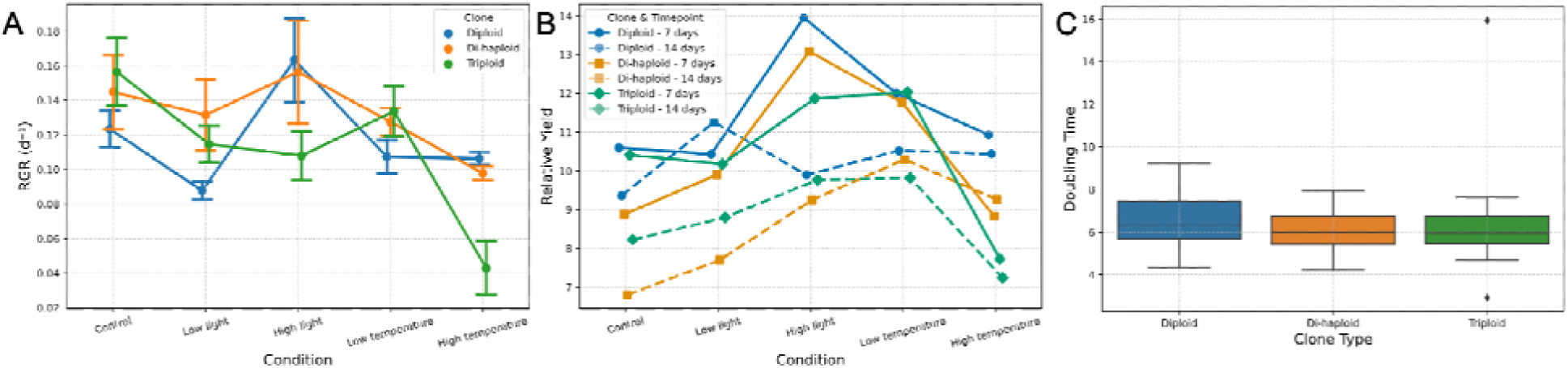
Growth characteristics of Lemna clones under varying environmental conditions. This figure illustrates the growth characteristics of *L. minor* diploid, *L. × japonica* di-haploid and *L. × japonica* triploid clones under different environmental conditions (see figure 6). A) Relative Growth Rate (RGR): RGR is shown for each clone type (diploid, di-haploid, and triploid) across five conditions: Control, Low light, High light, Low temperature, and High temperature. Error bars represent standard deviation. B) Relative Yield (RY): Comparison of RY between 7 and 14 days across the same environmental conditions. Different ploidy levels are indicated by different colors and the different time points (7 days vs. 14 days) by solid versus dashed lines. C) Doubling Time (DT): Boxplot showing DT distribution for each clone type.

